# Blood-brain barrier transport using a high-affinity, brain-selective VNAR (Variable Domain of New Antigen Receptor) antibody targeting transferrin receptor 1

**DOI:** 10.1101/816900

**Authors:** Pawel Stocki, Jaroslaw Szary, Charlotte LM Rasmussen, Mykhaylo Demydchuk, Leandra Northall, Diana Bahu Logan, Aziz Gauhar, Laura Thei, Torben Moos, Frank S Walsh, J Lynn Rutkowski

## Abstract

Transfer across the blood-brain barrier (BBB) remains a significant hurdle for the development of biopharmaceuticals with therapeutic effects within the central nervous system. We established a functional selection method to identify high-affinity single domain antibodies to the transferrin receptor 1 (TfR1) with efficient biotherapeutic delivery across the BBB.

**Methods:** A synthetic phage display library based on the variable domain of new antigen receptor (VNAR) was used for *in vitro* selection against recombinant human TfR1 ectodomain (rh-TfR1-ECD) followed by *in vivo* selection in mouse for brain parenchyma penetrating antibodies. Phage formatted VNARs cross-reactive to recombinant human and mouse TfR1-ECD were fused to Fc domain of human IgG1 (hFc) and tested for TfR1-ECD binding by ELISA and surface plasmon resonance. The pharmacokinetics and biodistribution of VNAR-hFcs were studied in mice by ELISA and immunolabeling following intravenous (IV) injection and cardiac perfusion. Functional activity was measured by body temperature reduction following the IV injection of neurotensin fused to a TXB2-hFc (TXB2-hFc-NT).

**Results:** TXB2 was identified as a high-affinity, species cross-reactive VNAR antibody against TfR1-ECD, that does not to compete with transferrin or ferritin for receptor binding. IV dosing of TXB2-hFc at 25 nmol/kg (1.875 mg/kg) in mice resulted in rapid binding to brain capillaries with subsequent transport into the brain parenchyma and specific uptake into TfR1-positive neurons. Likewise, IV dosing of TXB2-hFc-NT at 25 nmol/kg resulted in a rapid and reversible pharmacological response as measured by body temperature reduction. TXB2-hFc did not elicit any acute adverse reactions, bind or deplete circulating reticulocytes or reduce BBB-expressed endogenous TfR1 in mice. There was no evidence of target-mediated clearance or accumulation in peripheral organs except lung.

**Conclusions:** A species cross-reactive and brain-selective VNAR antibody to TfR1 was identified by a combination of *in vitro* and *in vivo* phage selection. As a high-affinity, bivalent Fc fusion protein, TXB2 rapidly crossed the BBB and exhibited a favorable pharmacokinetic and safety profile and can be readily adapted to carry a wide variety of biotherapeutics from blood to brain.

## INTRODUCTION

The blood-brain barrier (BBB) is formed by highly specialized endothelial cells that maintain an optimal environment for neuronal function by eliminating toxic substances and supplying the brain with nutrients and other metabolic requirements. While the BBB also poses a major challenge for drug delivery to the central nervous system (CNS), receptor-mediated transport systems for endogenous ligands have been exploited to shuttle a wide range of biologics into the brain in a non-invasive manner. Transferrin receptor 1 (TfR1), which mediates uptake of iron loaded transferrin (Tf) with subsequent transfer of iron into the brain and the return of iron-depleted Tf to the blood, is the most extensively characterized [1]. The OX26 monoclonal antibody, which binds the rat TfR1 without blocking the binding of Tf, was the first antibody exploited to carry a drug across the BBB [2]. Since then, the delivery of many biomolecules to the brain have been attempted using a variety of antibodies targeting the TfR1 [3].

Despite clear advances, several features of monoclonal anti-TfR1 antibodies as BBB carriers have hampered their clinical development. Anti-TfR1 antibodies can cause anemia by target-mediated lysis of TfR1-rich reticulocytes [4, 5]. Off-target binding to TfR1 expressed in other peripheral tissues such as liver and kidney, likely accounts for the relatively short plasma half-life of some antibodies targeting TfR1 [4, 6, 7]. Anti-TfR1 antibodies can accumulate in brain capillaries with limited release into the brain parenchyma [8] and can direct the receptor for lysosomal degradation [9, 10]. These antibodies have been subsequently re-engineered to reduce their safety liabilities and improve brain penetration either by monovalent formatting or a reduction in TfR1 binding affinity [8, 11-13]. However, these modifications necessitate higher doses to achieve a significant degree of receptor occupancy, and it is not yet clear whether CNS exposure of anti-TfR1 antibodies has been fully maximized. Moreover, anti-TfR1 antibodies used to date are species-specific, which remains problematic for preclinical efficacy and safety testing of potential therapeutic molecules.

We hypothesized that the variable domain of new antigen receptors (VNARs) derived from single domain antibodies found in the shark [14] could circumvent some of the drawbacks of the anti-TfR1 antibodies. The unique antigen binding paratope of VNARs harbors a long complementarity-determining region 3 (CDR3), which provides access to cryptic epitopes often not accessible by conventional monoclonal antibodies. Moreover, the small size, high solubility, thermal stability and refolding capacity of VNARs enable their multi-modal coupling to biopharmaceuticals [15].

In the current study, the VNAR TXB2 was identified from a semi-synthetic VNAR phage display library by a combination of *in vitro* and *in vivo* selections. We show that TXB2 specifically interacts with recombinant mouse, rat, monkey and human TfR1-ECD with high affinity (≤1 nM KD) without blocking the binding of Tf or ferritin to the transferrin receptor. In mice, TXB2 rapidly crosses the BBB as a bivalent VNAR-human Fc fusion protein (TXB2-hFc), and is readily detected in brain capillaries, parenchyma and neurons at low therapeutic dose (1.875 mg/kg). Neuronal delivery of a biologically active peptide following BBB transport was confirmed at the same dose with TXB2-hFc fused to neurotensin (NT) when injected intravenously. Additional biodistribution, pharmacokinetic and safety studies indicate that the high affinity TXB2 antibody has major advantages as BBB carrier.

## METHODS

### *In vitro* phage selections

Two semi-synthetic libraries based on type I and II nurse shark VNAR isoforms were designed to exclude human T-cell epitopes [16]. Phage libraries were produced in super-infected ER2738 cultures and PEG/NaCl precipitated. Two rounds of *in vitro* selections were performed on rh-TfR1-ECD (Sino Biological) that was biotinylated using Sulfo-NHS-Biotin EZ-Link kit (Thermo Fisher). The libraries were first deselected on streptavidin-coupled Dynabeads (Thermo Fisher) then used to pulldown phages bound to rh-TfR1-ECD. The beads were washed, eluted in 100 nM triethylamine and used for infection of *E. coli* strain ER2738 after adjusting to neutral pH. The output titer was calculated by counting antibiotic-resistant colonies, and the culture super-infected with M13KO7 helper phage to produce phages for subsequent rounds of selection.

### Phage ELISA

Individual clones were picked from agar plates and grown at 37°C with shaking in a 96-well block in 2YT media supplemented with 2% glucose, 5 μg/ml tetracycline and 100 μg/ml ampicillin until visible growth occurred. M13KO7 helper phage was then added and the media was replaced after 1 hr with 2YT media supplemented with 5 μg/ml tetracycline, 100 μg/ml ampicillin and 50 ug/ml kanamycin and grown for 16 hr at 30°C. Plates were then spun down at 2,500xg, and the supernatant recovered and blocked with final concentration of 2.5% milk in PBS with 0.1% Tween20 (PBST) for 15 min at room temperature (RT) before preceding to ELISA. Nunc MaxiSorp plates (Thermo Fisher) were coated with 100 µl of 1 µg/ml of recombinant mouse and human TfR1-ECD (Sino Biological) and human serum albumin (HSA; Sigma) and incubated at 4°C overnight. Plates were incubated with blocking buffer (2.5% non-fat dry milk in PBST) for 1 hr at RT. Blocked supernatants from the phage rescue were transferred to the blocked plates and incubated for 1 hr. The plates were washed with PBST and incubated with anti-M13-HRP 1:3,000 (GE Healthcare) in blocking buffer for 30 min. The plates were washed and developed with SureBlue™ TMB substrate (Thermo Fisher); the reaction was stopped with 1% HCl and absorbance measured at 450 nm.

### *In vivo* phage selections

Endotoxins were removed from phage preparations with Triton X-114 (final concentration of 1%). After incubated for 5 min on ice followed by 5 min at 37°C, preparations were centrifuged at 13,000 rpm for 5 min at 37°C, the supernatant was transferred to a new tube. The process was repeated twice, and endotoxin levels were measured using the LAL assay (Associates of Cape Cod, Inc.). Female BALB/c mice 6-12 weeks old received a single tail injection of a 100 µl phage preparation (5×10^11^ cfu in PBS) followed by perfusion with 25 ml of PBS supplemented with 1 EU/ml of heparin at either 1, 3 and 18 hr after injection.

### Brain fractionation and phage rescue

Brains were removed and the capillaries depleted from the parenchyma as previously described [17]. Briefly, brain tissue was homogenized with 14 strokes in a Dounce tissue grinder on ice in a 3:1 v/w of physiologic buffer (10 mM HEPES, 141 mM NaCI, 4 mM KCI, 2.8 mM CaCI2, 1 mM MgSO4, 1 mM NaH2PO4 and 10 mM D-glucose, pH 7.4), and the capillaries depleted by centrifugation through 13% dextran (60-90 kDa) at 5,400xg for 15 min at 4°C. A 1 ml aliquot of supernatant was added to 30 ml of *E. coli* ER2738 cells at mid-log phase growth in 2YT with tetracycline (5 µg/ml). The bacteria were further grown for 2 hr at 37°C while shaken at 250 rpm. After centrifugation at 3,000 rpm for 15 min, the bacterial pellet was plated onto 2YT agar plates containing ampicillin (100 µg/ml) and glucose (2% w/v), and incubated overnight at 30°C. Antibiotic-resistant colonies were counted, and a suspension culture was initiated from the agar plates and allowed to grow to mid-log phase in 2YT media with ampicillin (100 µg/ml) and glucose (2% w/v) before being super-infected with M13KO7 helper phage. The culture was further grown for 1 hr before the media was changed to 2YT with ampicillin (100 µg/ml) and kanamycin (50 µg/ml) and subsequently grown overnight at 30°C, while shaken at 250 rpm. The rescued phages were precipitated with PEG/NaCl, and endotoxins were removed before the next round of *in vivo* selection.

### Next generation sequencing

Phagemid DNA samples from each round of selection were prepared using QIAfilter kit (Qiagen). An approximately 350bp region between two SfiI restriction sites was sequenced using Illumina MiSeq platform in a 2×250 bp paired-end configuration (GeneWiz). An average Q score of greater than Q30 was obtained for 75% of the sequencing data where Q score refers to log-transformed error probability. Post-acquisition analysis included grouping VNARs into families with common CDR3s. Relative abundance was calculated per million sequences for each stage of the selection using RStudio analysis software.

### Production of human Fc and NT fusion proteins

VNAR-hFc formats were produced with VNARs at N-terminal end of hFc IgG1. Unless specified otherwise, hFc domain that was used for all the constructs throughout the study carried a series of mutations (E233P/L234V/L235A/ΔG236 + A327G/A330S/P331S) to attenuate effector function (AEF) [18]. The 8D3-hFc antibody was engineered with VH and VL domains of a rat anti-mouse specific TfR1 antibody [19] cloned into human heavy chain IgG1 with and human kappa light chain, respectively. NT constructs were designed to include the peptide on the C-terminal end of hFc following a 3xG4S linker. The Exp293F expression system (Thermo Fisher) was used for protein production following the manufacturer’s manual. After 5 days growth, the cells were centrifuged at 2,000 rpm for 10 min. Supernatants were filtered using 0.22 µm membrane filters and loaded onto HiTrap MabSelect SuRe (GE Healthcare) column pre-equilibrated against PBS, pH 7.4. Protein A affinity bound proteins were eluted with 0.1 M Glycine, pH 3.5 and the buffer exchanged to PBS, pH 7.4 using HiPrep 26/10 Desalting (GE Healthcare). Purity of the purified protein samples was determined by analytical size exclusion chromatography (SEC) and, if required, protein samples were concentrated and further purified by preparative SEC using a HiLoad 26/600 Superdex 200 pg (GE Healthcare) column. Purified proteins of interest were concentrated, filtered using 0.2 µm membrane filters and the quality of the final samples was confirmed by analytical SEC and sodium dodecyl sulfate polyacrylamide gel electrophoresis (SDS-PAGE). The integrity of the NT constructs was confirmed by mass spectrometry.

### Tissue ELISA

MaxiSorp plates were coated with 100 µl of goat anti-human Fc antibody (Sigma) diluted 1:500 in PBS or 3 mg/ml of an anti-VNAR antibody (courtesy of Martin Flajnik, University of Maryland, Baltimore) overnight at 4°C. Plates were washed and incubated with blocking buffer for 1 hr at RT. Tissue samples from brain and peripheral organs were homogenized in 3:1 (v/w) of PBS containing 1% Triton X-100 supplemented with protease inhibitors (cOmplete™, Sigma) using the TissueRuptor (Qiagen) at medium speed for 10 sec and then incubated for 30 min on ice. Lysates were spun down at 17,000xg for 20 min; the supernatant was collected and blocked in 2.5% milk in PBST overnight at 4°C. Blocked brain lysates (100 µl) were added to the blocked plates and incubated for 1 hr at RT. Plates were washed, and the VNAR-hFc concentration determined as described for the binding ELISA.

### TfR1 binding ELISA

Nunc MaxiSorp plates (Thermo Fisher) were coated with 100 µl of 1 µg/ml of recombinant mouse TfR1 (rm-TfR1-ECD; Sino Biological), rh-TfR1-ECD, HSA (Sigma), mouse transferrin receptor 2 ectodomain (TfR2-ECD; ACRO Biosystems) or in-house purified recombinant human, mouse, rat, monkey TfR1-ECDs and incubated at 4°C overnight. Plates were incubated with blocking buffer (2.5% non-fat dry milk in PBST) for 1 hr at RT. Supernatants from transfected cells or purified proteins were mixed with non-fat dry milk in PBST to a final concentration of 2.5% and incubated for 30 min. Blocked protein solutions (100 µl) were transferred to the blocked plates and incubated for 1 hr. Plates were washed with PBST and incubated with a goat anti-human Fc−peroxidase antibody diluted 1:5,000 (Sigma) in blocking buffer for 30 min. Plates were washed and developed with SureBlue™ TMB substrate, the reaction stopped with 1% HCl and absorbance measured at 450 nm. The VNAR-Fc concentration was determined from standard curves prepared individually for each fusion protein.

### Cell transfection

Exp293F cells (Thermo Fisher) were transiently transfected with pCMV3 plasmids with cloned untagged, full length TfR1 from either mouse (NM_011638.4; Sino Biological MG50741-UT), monkey (NM_001257303.1; Sino Biological CG90253-UT), rat (NM_022712.1; Sino Biological RG80098-UT) or human TfR1 (NM_001128148.1; Sino Biological HG11020-UT) using ExpiFectamine (Thermo Fisher) following the manufacturer’s instructions. The cells were grown in a shaking incubator at 350 rpm, 37°C with 8% CO_2_ for total of 2 days. The transfected cells were spun down at 2,000 rpm for 10 min and analyzed by flow cytometry.

### Flow cytometry

The cells were collected and 2×10^5^ cells/well were transferred into V-bottom 96-well plates. Cells were blocked with PBS containing 1% BSA (FACS buffer) for 10 min on ice and then washed and incubated with TXB2-hFc or control VNAR-hFc and either mouse Tf-Alexa Fluor 647 (Jackson ImmunoResearch) or human Tf-Alexa Flour 647 (Invitrogen) for 30 min on ice. The cells were washed and incubated on ice with secondary antibody anti-hFc-PE (eBioscience) for 20 min in the dark. The cells then were washed and resuspended in 200 µl of FACS buffer, and 3 µl of propidium iodide (1 mg/ml) added per well to assess dead cells in the analysis, which was performed on a CytoFlex flow cytometer (Beckman Coulter).

### Competition ELISAs

For competition with human holo-Tf, MaxiSorp plates were coated with 100 µl of rh-TfR1-ECD (Sino Biological) at the concentration of 5 µg/ml overnight at 4°C. Plates were washed with PBST and blocked for 1 hr with 2% BSA in PBST. In one version of the assay, control wells were left in blocking buffer whereas competition wells were incubated with 2.5 µM holo-hTf (Sigma) for 1 hr and washed before adding 100 µl of TXB2-hFc at the concentration ranging from pM to µM and incubated for 1 hr. In another variation, biotinylated hTf (bio-hTf; Sigma) at concentrations ranging from pM to µM in 0.1% BSA in PBST was added to the wells. After a 1 hr incubation, 100 µl of TXB2-hFc or holo-hTf at the concentration of 2.44 nM was added and further incubated for 1 hr. Following washing, 100 µl of either a 1:5,000 peroxidase-conjugated anti-human Fc antibody or 1:20,000 streptavidin Poly-HRP40 Conjugate (Fitzgerald) diluted in 0.5% BSA in PBST was added and incubated for 1 hr. The plates were washed, and absorbance was measured as for the binding ELISA.

To evaluate competition with ferritin, plates were coated with 100 μl of either 1 μg/ml of rh-TfR1-ECD (purified in-house) or liver purified ferritin (Bio-Rad, 4420-4804) overnight at 4°C. The plates were then washed with PBST and blocked for 1 hour with 300 μl of 2.5% BSA in PBST. For plates coated with rh-TfR1-ECD, VNAR-hFcs were added at concentrations ranging from μM to pM, either alone or in the presence of 100 nM ferritin, and incubated for 1 hr. For the ferritin-coated plates, 50 μl of rh-TfR1-ECD at 10 μg/ml was added for 1 hr. After washing, VNAR-hFcs were added at μM to pM concentrations and incubated for 1 hr. The plates were then washed and incubated with a 1:5,000 dilution of peroxidase-conjugated anti-human Fc antibody (Sigma) for 1 hr. After washing, absorbance was measured as for the binding ELISA.

### Binding kinetics

Binding kinetics of VNAR-Fcs was determined by surface plasmon resonance (SPR) using a Biacore T200 (GE Healthcare). A His-capture kit (GE Healthcare) was used to immobilize anti-His antibodies on CM5 chips (as recommended by the manufacturer). His-tagged recombinant ECD of mouse (NP_001344227.1; 123-763aa), rat (NP_073203.1; 123-761aa), monkey (XP_005545315.1; 121-760aa) and human (NP_001121620.1; 120-760aa) TfR1 in 0.1% BSA in HBS-EP+ buffer (GE Healthcare) was captured at flow rate 10 μl/min. Capture levels for mouse, rat, monkey and human TfR1-ECD were approximately 90, 47, 208 and 250 RU, respectively. The percentage of theoretical Rmax on average was 12, 26, 12, 58, 28 and 51% for control VNAR-hFc, TXB2-hFc, holo-hTf, OKT9, 8D3-hFc and OX26, respectively. Analyte binding was measured using the single cycle kinetic SPR method. Analytes including TXB2-hFc, 8D3-hFc, OX26 (Novus Biologicals), OKT9 (Invitrogen) and holo-hTf were injected at increasing concentrations (0.98, 3.9, 15.6, 62.5 and 250 nM) in HBS-EP+ at flow rate 30 µl/min. A flow cell without TfR1 captured served as a reference. The chips were regenerated in 10 mM Glycine-HCl, pH 1.5. Sensorgrams were fitted using 1:1 binding model and kinetic constants determined using Biacore T200 Evaluation software (GE Healthcare).

### Reticulocyte binding and depletion

Antibody binding to mouse reticulocytes was assessed *ex vivo* using fresh blood samples diluted 1:5 with PBS. Aliquots (10 µl) were added to 1 ml of BD Retic-Count (BD Biosciences) for 60 min. The samples were then stained for 30 min at RT in the dark with molar equivalents of either TXB2-hFc or 8D3-hFc (10 or 20 µg/ml, respectively) followed by 30 min incubation with 20 ug/ml of anti-hFc antibody conjugated to Alexa Flour 647 (Thermo Fisher). Antibody binding to the reticulocyte population was determined by flow cytometry (CytoFlex). To assess the effect of antibody exposure on circulating reticulocytes *in vivo*, mice were injected IV with TXB2-hFc or 8D3-hFc at doses of 25 and 250 nmol/kg, and blood was collected after 18 hr. PBS injected animals were used as a control. Blood samples were diluted 1:5 with PBS and 10 µl aliquots were added to 1 ml of BD Retic-Count reagent (BD Biosciences). After a 60 min incubation in the dark at RT, the samples were analyzed by flow cytometry.

### Western blotting

To assess changes in TfR1 expression after antibody exposure, whole brain lysates were prepared as for ELISA and protein concentration determined using a BCA assay (Thermo Fisher). Wells were loaded with 30 µg of total protein which was resolved by SDS-PAGE under reducing conditions and transferred onto polyvinylidene difluoride (PVDF) membranes (Thermo Fisher). After blocking with 2% BSA for 30 min, the membranes were incubated with mouse anti-TfR1 clone H68.4 (Thermo Fisher) and rabbit anti-actin antibodies (Abcam). Binding was detected with an anti-mouse Cy-3 and anti-rabbit Cy-5 antibodies (GE Healthcare). Blots were scanned using a Typhoon 5 (GE Healthcare) and the signal intensity quantified using Image Studio Lite (LI-COR Biosciences).

To monitor the efficiency of capillary depletion, protein concentrations in capillary and parenchymal fractions were measured using the BCA assay. Wells were loaded with 10 µg of total protein in reducing buffer without boiling. The protein samples were resolved by SDS-PAGE and transferred to PVDF membrane and blocking as above before incubation with rabbit anti-GLUT1 (Abcam) and mouse anti-tubulin (Abcam) antibodies. The detection and quantitation were performed as for TfR1 above.

### Immunohistochemistry (IHC)

To assess brain penetration of TXB2-hFc *in vivo*, 20 female BALB/c mice (6 wk old), were acclimatized for 14 days prior to treatment (approved under license no: 2018-15-0201-01550 by the Danish Experimental Animal Inspectorate under the Ministry of Food and Agriculture). Mice received a tail vein injection with either control VNAR-hFc or TXB2-hFc in doses of 2 or 12 mg/kg (27 nmol/kg and 160 nmol/kg, respectively). After a 1 or 18 hr exposure, mice were deeply anesthetized with a subcutaneous injection of 0.1 ml/10 g of body weight with fentanyl (0.315 mg/ml), fluanisone (10 mg/ml), midazolam (5 mg/ml) diluted in sterile water (1:1:2) and perfused transcardially with 0.01M potassium phosphate-buffered saline (PPBS) followed by 4% paraformaldehyde (PFA) in PPBS, pH 7.4. Brains were dissected and post-fixed in 4% PFA at 4°C overnight, extensively washed in PPBS, and then transferred to 30% sucrose for a minimum of 48 hrs.

Serial coronal brain sections (40 µm) were cut on a cryostat (Leica CM3050 S), and collected free-floating in PPBS, pH 7.4, in a sequential series of six and stored at -20°C in anti-freeze solution. For staining, free-floating brain sections were pre-incubated in blocking buffer (3% porcine serum diluted in 0.01 M PPBS with 0.3 % Triton X-100) for 60 min at RT and then incubated overnight at 4°C with biotinylated goat anti-human IgG antibody (Vector, BA 3000) diluted 1:200 in blocking buffer. After 24 hr, the sections were washed twice in washing buffer (blocking buffer diluted 1:50 in PPBS) and once in PPBS. Antibodies were visualized using the Avidin–Biotin Complex-system (ABC-HRP kit, Vectastain) and 3,3′-diaminobenzidine tetrahydrochloride as the substrate.

*Ex vivo* binding of TXB2-hFc was evaluated in brains taken from adult female BALB/c mice that were fixed in 4% PFA at 4°C overnight, extensively washed in PPBS and paraffin embedded. Serial sections (6 µm) were treated with xylene, dehydrated with ethanol and exposed to 3% hydrogen peroxide for 10 min to inactivate endogenous peroxidases. Antigen was retrieved by boiling in 10 mM citrate buffer pH6 for 10 min. Sections were blocked with 3% porcine serum albumin in Tris-buffered saline (TBS) for 1 hr at RT and incubated with primary antibodies at 100 μg/ml overnight at 4°C. Sections were washed in TBS then incubated for 1 hr at RT with a biotinylated goat anti-human IgG (H+L) antibody (Vector BA3000; 1:500 dilution). Antibody binding was detected using the Avidin– Biotin Complex-system (ABC-HRP kit, Vectastain) and 3,3′-diaminobenzidine tetrahydrochloride as the substrate. All images were captured on a brightfield Zeiss Axioskop microscope at 20X magnification with the same acquisition parameters.

### Immunofluorescent labelling

For double-fluorescent labeling of TXB2-hFc and endogenous TfR1, sections were co-incubated overnight at 4°C with biotinylated goat anti-human IgG antibody and rat anti-mouse TfR1 (clone RI7217, synthesized by Aalborg University) diluted 1:200 in blocking buffer. Sections were then washed and incubated with Alexa Fluor 594-conjugated donkey anti-rat IgG diluted 1:200 for 30 min (for TfR1) and successively with the Avidin–Biotin Complex-system for 30 min, biotinylated-tyramide (diluted 1:100 for 5 min), Avidin–Biotin Complex-system for 30 min, and finally streptavidin-conjugated Alexa Fluor 488 to detect TXB2-hFc. To detect the presence of TXB2-hFc in glial cells, sections were co-incubated with polyclonal rabbit anti-GFAP (Dako) for astrocytes, polyclonal rabbit anti-P25 alpha (courtesy of Poul Henning Jensen, Aarhus University) for oligodendrocytes, or monoclonal rat anti-IBA1 (Serotec) for microglia. The glial marker antibodies were all diluted 1:200 in blocking buffer and detected with donkey anti-rabbit IgG conjugated with Alexa Fluor 594 (1:200, Life Technologies) for 30 min at RT. The fluorescent sections were examined and imaged using the fluorescence DMi8 Confocal Laser Scanning Microscope (Leica).

### Body temperature measurements

Five mice per group received tail vein injections with either VNAR-hFc-NT fusions or control hFc-NT at dose from 2.5 to 125 nmol/kg IV (0.2 mg/kg and 9.4 mg/kg) and body temperature was measured over a 24 hr period using a rectal probe, and the data analyzed using Prism software.

## RESULTS

### VNAR phage display selections

Semi-synthetic VNAR libraries were first subjected to 2 rounds of panning on rh-TfR1-ECD followed by 3 rounds of *in vivo* selection in mice. A flow diagram of the entire selection and screening process is shown in **Fig. S1**. The *in vivo* stability of the preselected phage library was tested by injecting 5×10^11^ cfu and harvesting perfused brains 1, 3 and 18 hr later. Capillaries were depleted by density centrifugation and the parenchymal fraction was used to infect ER2738 E. coli. The phage output titer was determined and approximately 100 clones were randomly picked from each of 1- and 18-hr time points. Binding to rh- and rm-TfR1-ECD was assessed by phage ELISA. Analysis showed that phage titers in addition to the number of TfR1 binders decreased with time after injection (**Fig. S2A**); thus a 1-hr exposure was used for the subsequent 3 rounds of *in vivo* selections.

Next generation sequencing (NGS) was used to follow individual clones throughout the selection. Tracing the 29 most abundant clones after the last round of selection back through the entire process showed a consistent amplification at both the *in vitro* and *in vivo* stage (**Fig. 1A and Table S1**) [20]. However, when the 20 most abundant clones after the *in vitro* stage were tracked forward, only a portion of clones continued to amplify whereas most were de-selected *in vivo* (**Fig. S2B**). Additionally, phage ELISA showed that the number of TfR1-ECD binders increased with each round of *in vivo* selection starting from 5% after round 1 to 15% after round 3 (**Fig. S2C**). These results indicate that the *in vivo* selection process effectively amplified functionally relevant TfR1-ECD binders.

**Figure 1.**
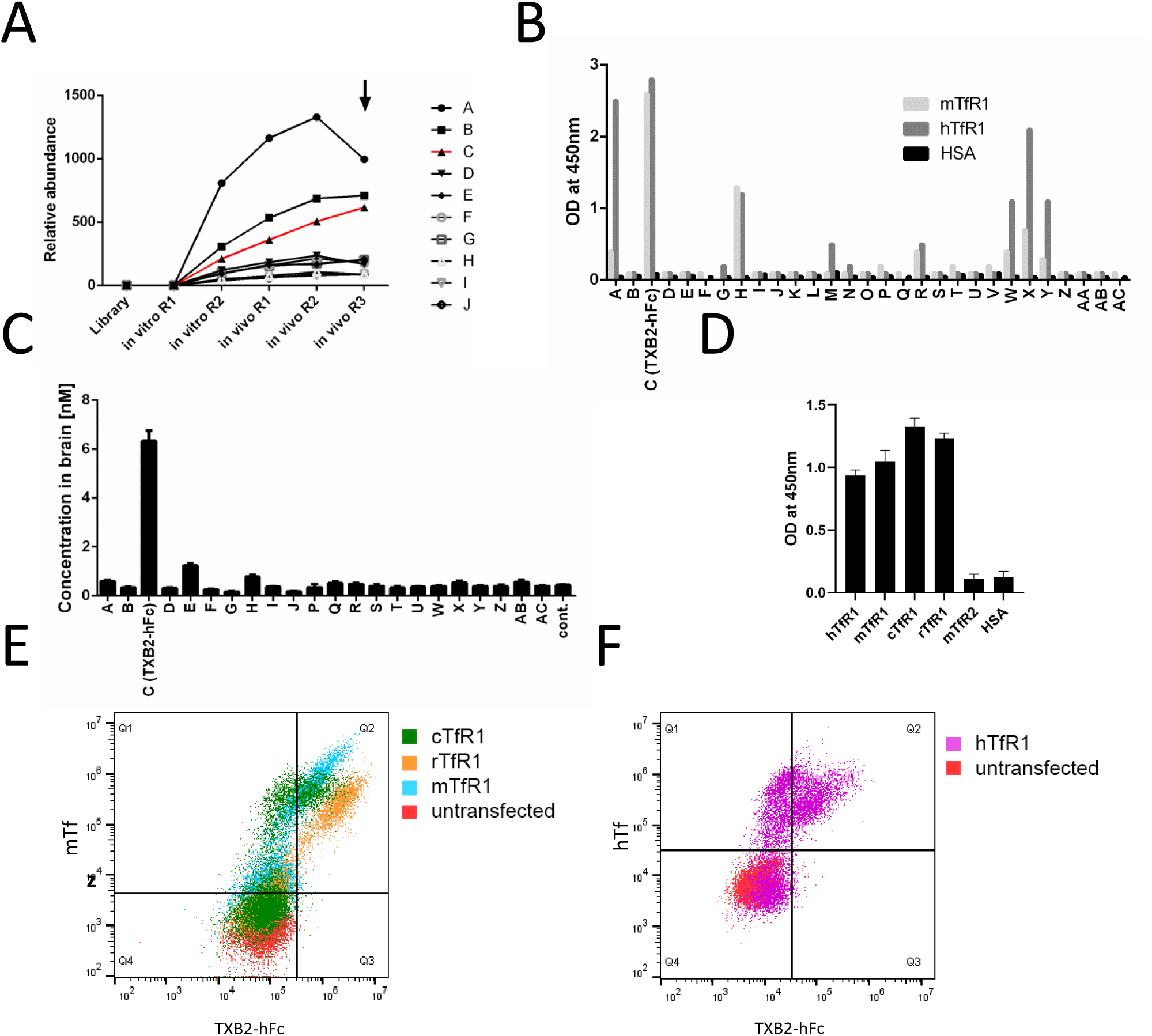
Selection, TfR1 binding and brain uptake of selected VNAR clones. (**A**) The selection process was monitored by NGS and the relative abundance of each VNAR family (CDR3-grouped) per million sequences was calculated. The progressive enrichment of the 10 most abundant clones (Table S1) from in vivo round 3 (marked by arrow) was represented in earlier rounds of selection. (**B**) Relative ELISA binding of the 29 most enriched VNARs (Table S1) formatted as hFc fusions to TfR1-EDCs of human and mouse (hTfR, mTfR1). (**C**) Brain concentration of the 22 tested VNAR-hFc fusions 18 hr after single IV injection of 1.875 mg/kg (25 nmol/kg) as determined by ELISA (mean ±SD, N=5). Control VNAR-hFc that binds to mouse TfR1 (Table S2, Figure S3) was included as negative control (cont.). (**D**) Comparative ELISA binding of TXB2-hFc to the ECDs of TfR1 of human, mouse, cynomolgus monkey and rat (hTfR, mTfR1, cTfR1, rTfR1), with mouse TfR2-ECD (mTfR2) and HSA as negative controls (mean ±SD, N=3). (**E**) Live HEK293 cells transiently transfected with mTfR1, cTfR1, rTfR1 or (**F**) hTfR1 were co-stained with TXB2-hFc and either transferrin from mouse (mTf) or human (hTf) and analyzed by flow cytometry. Double-positive cells that populated Q2 after transfection showed binding of both TXB2-hFc and Tf to the transiently expressed TfR1.

The 29 most abundant clones after the final round of *in vivo* selection (**Table S1**) were re-engineered as bivalent VNAR-hFc fusion proteins (**Fig. S3A**) to further evaluate their functional activity. Among the 29 clones that were tested, 8 showed TfR1-ECD binding by ELISA, and 7 were cross-reactive to the human and mouse receptor (**Fig. 1B**). The 22 clones that expressed well as VNAR-hFc fusions were tested for brain uptake in mice 18 hr after IV injection at 1.875 mg/kg (25 nmol/kg). A VNAR-hFc that binds TfR1-ECD with nM affinity (**Table S2**) but does not penetrate the BBB was used as a negative control (control VNAR-hFc) throughout the experiments. Brain uptake of TXB2-hFc (clone C) was 14-fold higher than the control and reached 6 nM in the brain 18 hr after injection (**Fig. 1C**). Identical results were obtained using either Fc capture or a VNAR capture ELISAs (**Fig. S3B**), which confirmed the integral stability of the fusion molecule upon brain delivery. Two additional clones (E and H) showed a modest increase in brain uptake over the control (4- and 2-fold, respectively), but only clone H bound rhTfR1-ECD.

### Binding properties

TXB2, which is a type II VNAR containing a disulfide bond between cysteine residues in CDR3 and CDR1, was selected for further characterization. Species cross-reactivity and specificity of TXB2-hFc was tested with the soluble, recombinant TfR1-ECD and with the full-length receptor expressed in a cell membrane. ELISAs showed TXB2-hFc binding to mouse, human, cynomolgus monkey and rat TfR1-ECD, while no binding was observed to HSA or mouse TfR2-ECD, the nearest homolog (**Fig. 1D**). The binding of TXB2-hFc to human, mouse, rat and cynomolgus monkey TfR1-ECDs was further analyzed by surface plasmon resonance (SPR). TXB2-hFc bound with sub-nM affinity to all but rat TfR1, which measured 1.2 nM KD and the binding kinetics were very similar to 8D3, OKT9 and OX26 antibodies, which are selective for mouse, human and rat TfR1, respectively (**Table S2**). Holo-hTf bound to hTfR1-ECD with a 6.8 nM KD in this setup. The control VNAR-hFc also showed strong nM binding to all but rat TfR1-ECD (**Table S2**) yet was unable to penetrate the BBB in mice.

TXB2-hFc binding to cell surfaces expressing TfR1s from various species was assessed in transiently transfected HEK293 cells by double staining for Tf and TXB2-hFc. Mouse Tf was used for cells transfected with mouse, rat and cynomolgus monkey TfR1 and human Tf for cells transfected with human TfR1. Flow cytometry showed that TXB2-hFc bound to full length TfR1 from all of the tested species when expressed on the surface of HEK293 cells (**Figs. 1E-F**). Binding of the control VNAR-hFc to mTfR1 expressed on the surface of HEK293 cells was also confirmed (**Fig. S4**).

A series of ELISAs were configured to test the competition between hTf and TXB2-hFc. Using plates coated with rh-TfR1-ECD and pre-incubation with biotinylated hTf (bio-Tf) at a constant concentration of 2.5 µM, there was no observed competition for TfR1 binding with TXB2-hFc at concentrations ranging from pM to µM (**Fig. S5A**). In the opposite configuration where TXB2-hFc was kept constant at 2.4 nM, increasing concentrations of bio-Tf used did not alter antibody binding to hTfR1 (**Fig. S5B**). In the same setup, when bio-Tf binding was challenged with unlabeled hTf, there was observable competition in binding to TfR1, but no competition was detected with a TXB2-hFc challenge (**Fig. S5C**). Thus, no competition between hTf and TXB2-hFc for binding to rh-TfR1-ECD was detected in these molecular assays.

TXB2-hFc was also tested for competition with ferritin, which competes with Tf for TfR1 binding [21]. VNAR-A06-hFc, which binds to rh-TfR1-ECD with a similar EC50 as TXB2-hFc (**Fig. S6A**) was used for comparison. In the first version of the assay, ELISA plates coated with ferritin were used to capture rh-TfR1-ECD before being incubated with VNAR-hFcs. The binding of TXB2-hFc to hTfR1 was unaffected whereas VNAR-A06-hFc did not bind hTfR1 that was pre-bound to ferritin (**Fig. S6B**). In the second version, ELISA plates coated with rh-TfR1-ECD were exposed to VNAR-Fcs in the presence of excess of ferritin. Again, VNAR-A06-hFc binding to hTfR1 was abolished, whereas TXB2-hFc binding remained unaffected by the presence of ferritin (**Fig. S6C**). In summary, these studies conclusively show that the high-affinity binding site for TXB2-hFc on rh-TfR1-ECD does not overlap with either Tf or ferritin.

### Brain penetration

To further examine BBB transport, brains were fractionated and capillary depletion from the parenchyma was confirmed by Western blotting for GLUT1 (**Fig. 2A**) as a specific marker for cerebral microvessels [22]. The amount of TXB2-hFc in the parenchyma was more than twice that in the capillary fraction by 18 hr after IV injection of 1.875 mg/kg (25 nmol/kg), whereas there was little of control VNAR-hFc in either fraction at the same dose (**Fig. 2B)**. In a longitudinal study, the concentration of TXB2-hFc in the capillary fraction was higher than the parenchyma 15 min after injection and then remained relatively constant over a 144 hr study (**Fig. 2C**). In contrast, the parenchymal concentration rose steadily over 18 hr and had only slightly declined by 144 hr. Results from IHC studies provided further evidence that TXB2-hFc penetrates the BBB into the brain parenchyma after IV injection. TXB2-hFc localized to the capillaries 1 hr after injection and was then visible in the adjacent parenchyma and cellular elements with neuronal morphologies by 18 hr (**Fig. 2D**). TXB2-hFc immunoreactivity within the capillary endothelial cells appeared independent of the dose (2 vs. 12 mg/kg, **Fig. 2D**), which suggested saturation of TfR1s at the BBB. Brain fractionation studies corroborate this result as capillary TXB2-hFc levels remained relatively constant throughout the 144 hr experiment (**Fig. 2C**).

**Figure 2.**
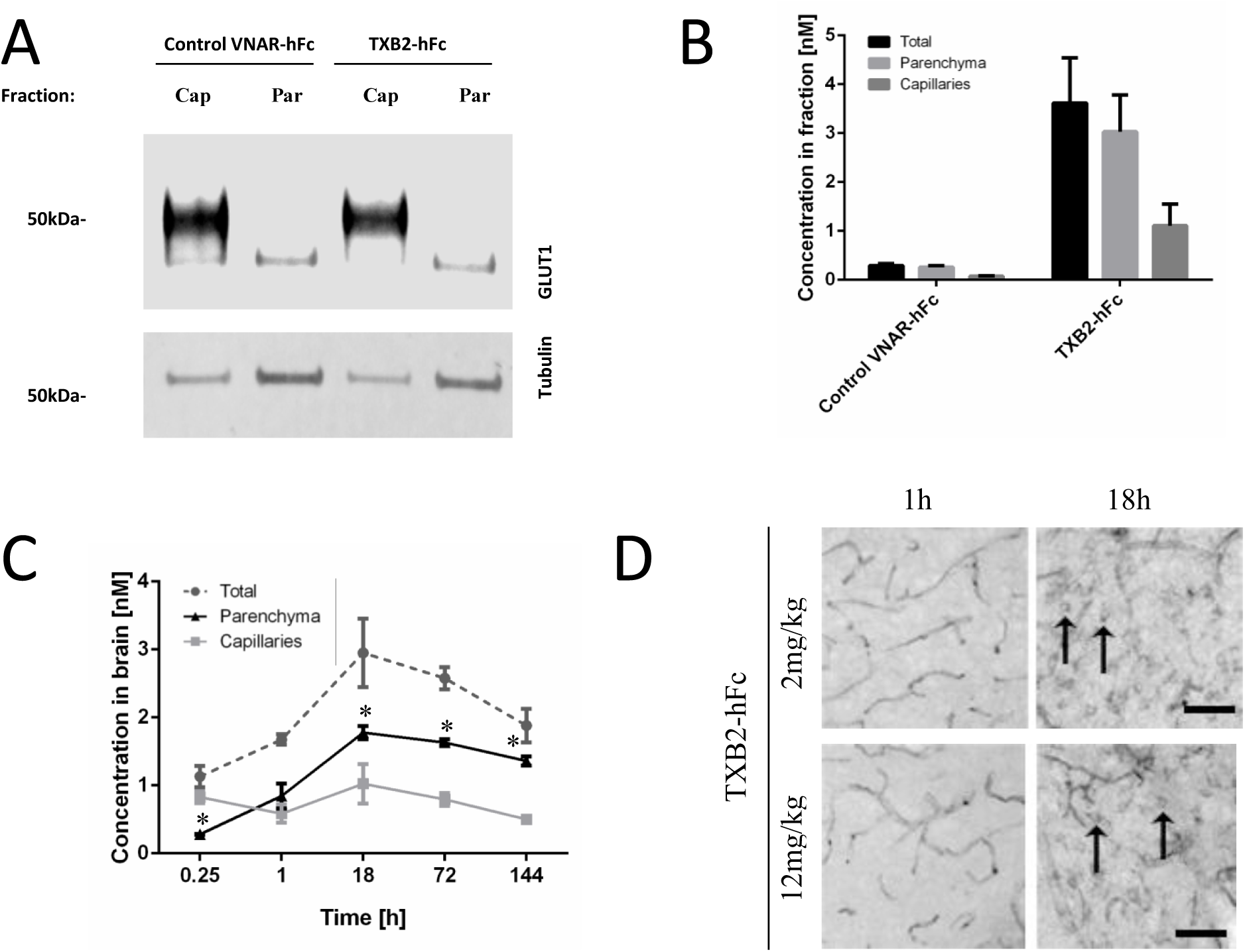
Parenchymal distribution of TXB2-hFc in brain after IV injection. (**A**) WB of capillary and parenchymal fractions. Separation of capillary (Cap) and parenchymal (Par) fractions after density centrifugation was shown by SDS-PAGE followed by WB for GLUT1 (capillaries) and tubulin (loading control). (**B**) Mice were administered TXB2-hFc or control VNAR-hFc at 1.875 mg/kg (25 nmol/kg) IV and VNAR-Fc concentrations in the capillary and parenchymal fractions compared to the unfractionated contralateral hemisphere (total) (mean ±SD, N=3). (**C**) Concentrations of TXB2-Fc in brain fractions over time after a single IV injection of 1.875 mg/kg (mean ±SD, N=3). * Indicates significant difference between parenchymal and capillary fractions (p< 0.05) by Student’s t-test. (**D**) Brain section from the superior colliculus of the mesencephalon. TXB2-hFc was injected at 2 or 12 mg/kg and allowed to circulate for 1 or 18 hrs. Immunolabeling was restricted to brain capillaries at 1 hr but after 18 hr additional parenchymal staining was observed; cells with a neuronal morphology were marked (arrows). Scale bar 50 µm.

### Brain distribution

TXB2-hFc immunoreactivity was observed in brain capillary endothelial cells, choroid plexus epithelial cells, and neurons after IV dosing (**Figs. 2D, 3 and S7**). Examining the surface of the brain revealed no significant signs of penetration across the pia-arachnoid barrier either after 1 hr (**Fig. S7C**) or 18 hr (**Figs. 3 A, E, H**). TXB2-hFc immunoreactivity was observed in epithelial cells of the choroid plexuses in the lateral, third (**Fig. S7B**), and fourth ventricles. Regions of the brain close to ventricular surfaces were carefully examined for evidence that TXB2-hFc may have crossed the BCSFB and entered the brain parenchyma by diffusion. In the diencephalon, faint labeling was seen in neurons of thalamic nuclei, but was unexpectedly absent from the medial habenular nucleus (**Fig. S7B**) known to strongly express TfR1 [23]. This finding might be attributed to the proximity to the ventricular surface and the flow of the brain interstitial fluid towards the ventricular compartment. Elsewhere in the brain, TXB2-hFc immunoreactivity closely mirrored the level of neuronal TfR1 expression (**Fig.3**), with no indication that neuronal labeling resulted from diffusion into the brain parenchyma from the CSF.

**Figure 3.**
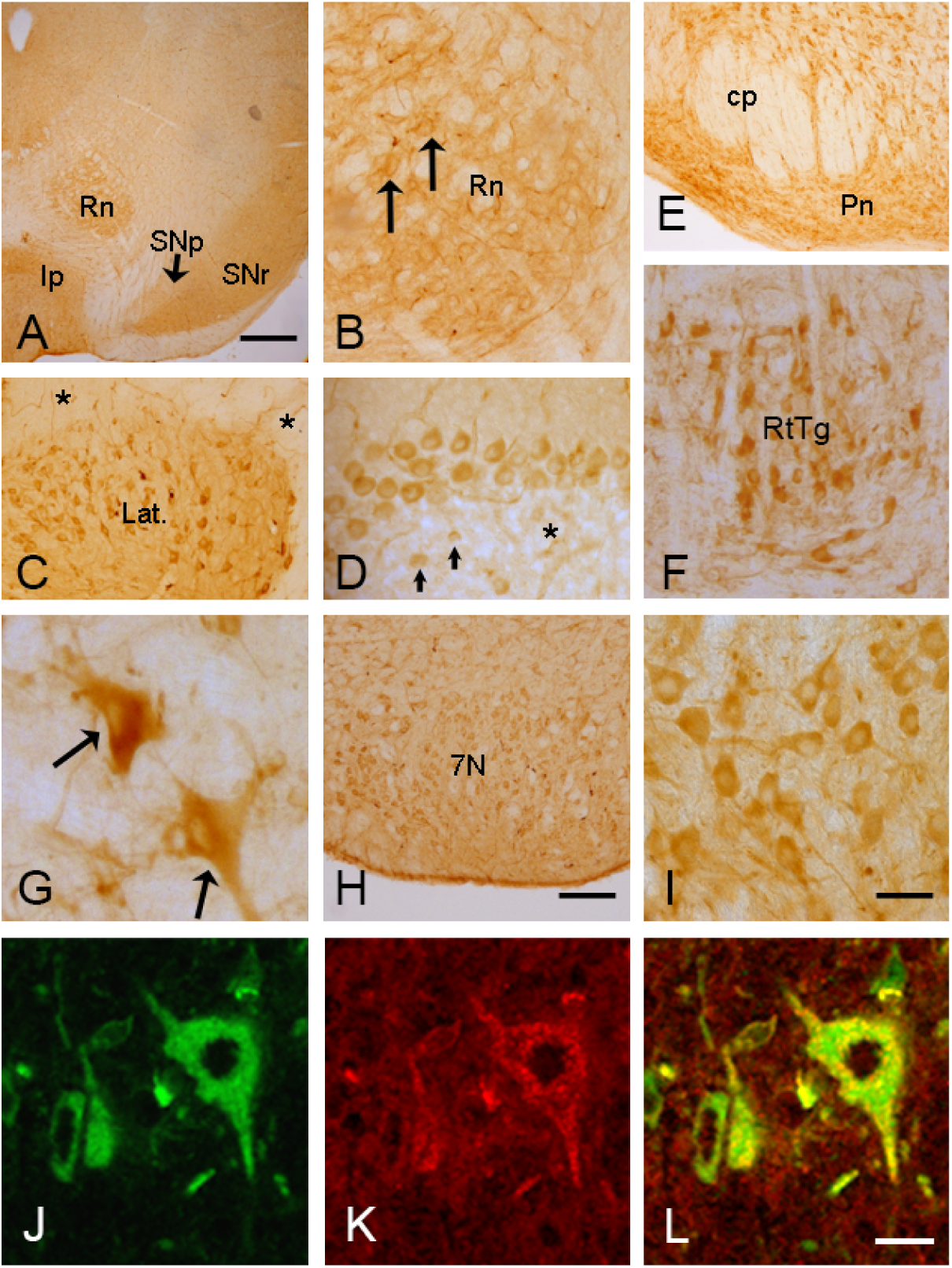
The distribution of TXB2-hFc in brain stem and cerebellum. IHC brain tissue analysis 18 hr after a dose of 12 mg/kg IV. The mesencephalon (**A**), with the red nucleus (Rn), interpedunculate nucleus (Ip), substantia nigra pars compacta (SNp) and substantia nigra pars reticulata (SNr). The Rn is shown in higher magnification (**B**) with neurons (arrows). The cerebellum (**C, D**), with the lateral nucleus (Lat.) and Purkinje cells (**D**), with discrete granular cells (arrows) and brain capillaries (*). The lower brain stem (**E-I**), with the reticular formation and cranial nerve nuclei. The pons (**E**) with the pontine nucleus (Pn), white matter-rich cerebral peduncle (cp) and neuronal labeling in the reticulotegmental nucleus (RtTg) (**F**). (**G**) Neuronal labeling is prominent in the reticular formation (arrows). (**H**) Labeling of neurons of the facial nucleus in lower and (**I**) higher magnification. Double fluorescent images taken in upper brain stem showing labeled neurons of the red nucleus. Neurons stained for (**J**) TXB2-hFc (green) and (**K**) TfR1 (red). (**L**) Merge image with green and red channels overlay showing double-labeling in yellow. Scale bars: **A** 1 mm, **B-D**,**F**,**I** 50 µm (bar shown in I), **E**,**H** 200 µm (bar shown in H), **G**,**J-L** 5 µm (bar shown in L).

Labeling with TXB2-hFc appeared stronger and more consistent in mice 18 hr after injection with 12 mg/kg, which more clearly illustrates neuronal uptake throughout the brain (**Fig. 3 A-I**). Labeling was highest in neurons of brain stem and cerebellum, but it was also detected in the diencephalon and telencephalic regions. In the neocortex, neuronal labeling was occasionally seen in cortical layers and pyramidal neurons of the hippocampus (not shown). In the mesencephalon (**Fig. 3 A, B**), TXB2-hFc immunoreactivity was discernable in reticular formation and cranial nerve nuclei and also seen markedly in neurons of the substantia nigra pars reticulata, more lightly in neurons of the substantia nigra pars compacta, and markedly in the interpedunculate nucleus. In the cerebellum (**Fig. 3 C, D**), TXB2-hFc immunoreactivity was high in Purkinje cells and neurons of deep cerebellar nuclei. In the brain stem (**Fig. 3 E-I**), TXB2-hFc immunoreactivity was prominent in high iron-containing regions and with neurons exhibiting strong TfR1 expression. Such regions include the substantia nigra pars reticulata, deep cerebellar nuclei of the cerebellum, and many neurons of the reticular formation [23, 24]. The neuronal localization of TXB2-hFc in brain stem regions extended from the medulla oblongata rostrally to the midbrain. In these regions, labeling was particularly high in motor cranial nerve nuclei and pontine nuclei. Brains sections examined for control VNAR-hFc showed no immunoreactivity in brain capillaries or neurons; however, labeling was seen in choroid plexuses of all ventricles suggesting non-specific transport of this VNAR at blood-CSF barrier (BCSFB). Faint immunoreactivity of control VNAR-hFc was confined to the innermost parts of the ventricular surfaces and the surfaces of the brain (**Fig. S7 D-F**) suggesting that BCSFB transport of control VNAR-hFc resulted in very limited diffusion into the brain parenchyma.

The biodistribution pattern of TXB2-hFc in the brain after IV administration closely mimicked the IHC staining pattern seen after direct antibody application to paraffin-embedded brain sections from untreated mice (**Fig. S9**). Like 8D3-hFc, TXB2-hFc showed intense staining of neurons of cortex layers II/III and in regions of large neuron-containing structures as well as those with a high density of axonal termini, including the CA2 of the hippocampus, the striosome bundles of the striatum, and the Purkinje and granular layers of the cerebellum. TXB2-hFc also reacted with localised blood vessels in all of these regions. By contrast, the irrelevant VNAR-G12 control as well as the control VNAR, which binds TfR1 under some conditions, did not show any selective staining of cellular or vascular elements.

### Neuronal specificity

Although TXB2-hFc immunoreactivity in the brain stem and cerebellum closely mirrored the level of neuronal TfR1 expression as published previously [23], double-fluorescent labeling unequivocally confirmed that TXB2-hFc co-localizes with TfR1-postive neurons (**Fig. 3 J-K**). By contrast, TXB2-hFc immunoreactivity was absent in the major glial cell types including astrocytes, microglia and oligodendrocytes (**Fig. S8**). Hence, in pure white matter regions such as the corpus callosum and cerebral peduncle, TXB2-hFc immunoreactivity was only detected in brain capillary endothelial cells. The immunolabeling of GFAP and IBA1 did not reveal any signs of reactive gliosis indicating that TXB2-hFc did not evoke inflammation in the brains when crossing the BBB.

### Biodistribution and safety assessment

For the *in vivo* biodistribution study, mice were perfused 18 hr after a single IV dose of 1.875 mg/kg (25 nmol/kg) of either TXB2-hFc, an irrelevant VNAR-G12-hFc control and the control VNAR-hFc, which bind TfR1 *in vitro* but not *in vivo*. The antibody concentrations in various organs were measured by ELISA **(Fig. 4A**). The mean plasma concentrations were similar for the three VNAR antibodies, indicative of similar PK profiles. While all the VNAR antibodies diffused into peripheral organs, only TXB2-hFc was found in the brain and achieved a concentration similar or greater than that in other tissues. The average molar concentration of TXB2-hFc in brain was 13- and 21-fold higher compared to VNAR-G12-hFc and control VNAR-hFc, respectively. The lung is the only peripheral organ where TXB2-hFc showed a significant increase of 1.6-fold when compared to control VNAR-hFc but not VNAR-G12-hFc.

**Figure 4.**
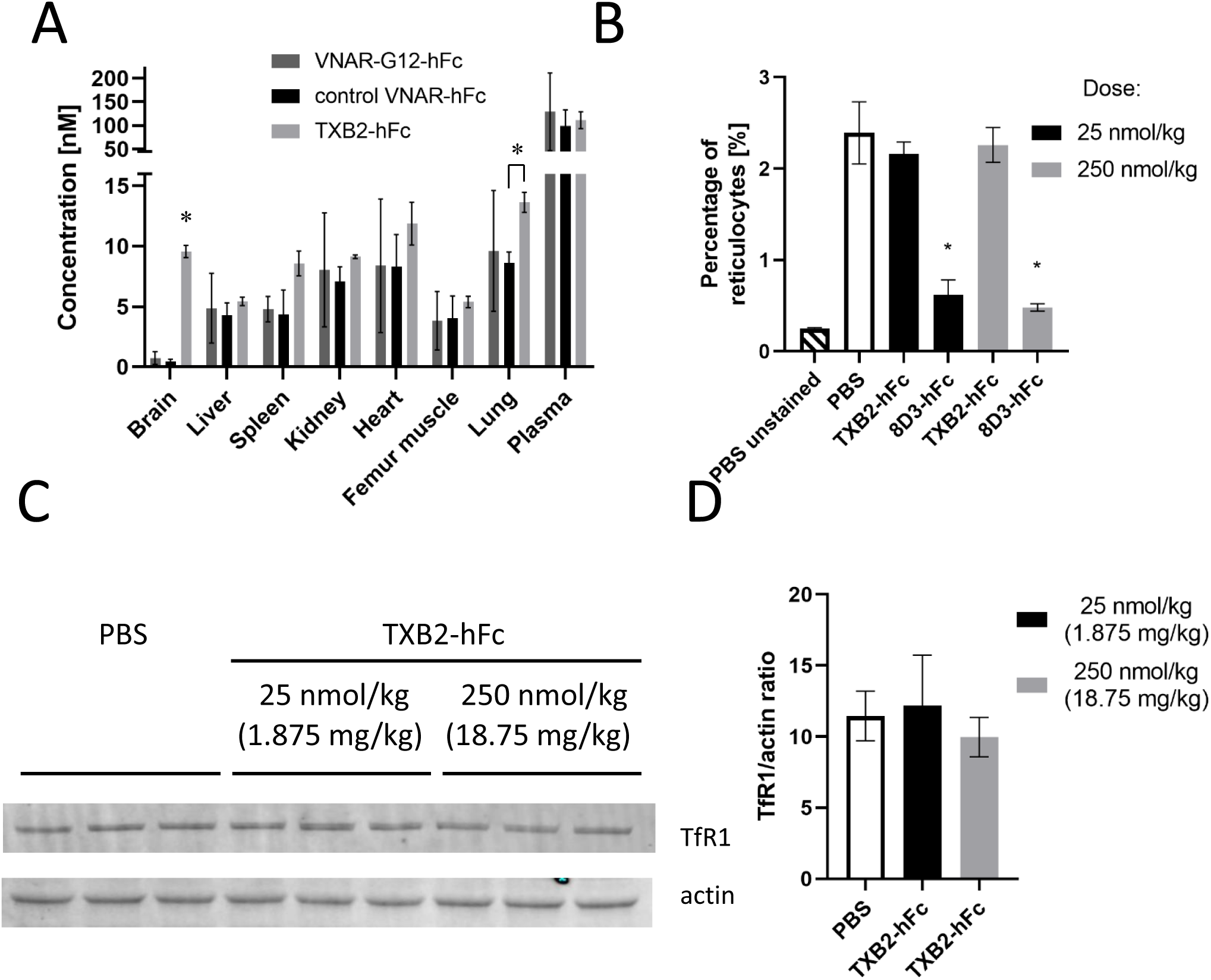
Biodistribution of TXB2-hFc and effect on circulating reticulocyte count or TfR1 expression in brain. (**A**) Biodistribution of TXB2-hFc compared to VNAR-G12-hFc (TfR1 non-binder), control VNAR-hFc (binds recombinant and cell expressed mouse TfR1), in mouse liver, spleen, lung, heart, kidney, femur muscle, brain and plasma. Mice were perfused and organs collected 18 hr after an IV injection of 1.875 mg/kg (25 nmol/kg). Tissues were homogenized and the antibody concentrations measured by ELISA (mean ±SD, N=3). * Indicates significance (p< 0.01) by Student’s t-test. (**B**) Mice were treated with TXB2-hFc or 8D3-hFc at 25 or 250 nmol/kg IV and PBS were used as a control. Blood samples were collected 18 hr post injection and the percentage of reticulocytes in total number of red cells (mean ±SD, N=3) was determined by flow cytometry after staining with Thiazole Orange. The unstained samples from PBS treated animals were used to setup the gate threshold at 0.25%. * Indicates significance (p< 0.05) by Student’s t-test when compared to PBS. (**C**)TXB2-hFc at a dose of 1.875 or 18.75 mg/kg (25 or 250 nmol/kg) was injected IV and perfused brains were harvested 18 hr later. Brain homogenates were analyzed by SDS-PAGE followed by WB using fluorescently labelled antibodies to detect TfR1 and actin. (**D**) WB data presented as a ratio of the TfR1 to actin signal (mean ±SD, N=5)

Serious adverse effects of TfR1 antibodies in mice as a consequence of targeting reticulocytes, which express high levels of TfR1 have been previously reported [4]. To assess this potential safety liability, a quantitative analysis of reticulocytes was performed in mice 18 hr after injecting TXB2-hFc or 8D3-hFc at 25 or 250 nmol/kg relative to vehicle controls. Both antibodies were fused to an Fc with attenuated effector function (AEF), which was shown to mitigate against rapid ADCC-mediated hemolysis [4]. 8D3-h Fc (AEF) still showed a significant reduction of reticulocytes even at the low dose of 25 nmol/kg (**Fig. 4B**). Increasing the dose by 10-fold did not exacerbated the effect, suggesting that the reduction was maximal at the lower dose. This residual level of toxicity was likely due to activation of the complement system as previously shown for a different TfR1 antibody with the same effector mutations [4]. By contrast, no reticulocyte depletion was seen with TXB2-hFc at either dose (**Fig. 4B**). Furthermore, TXB2-hFc also had no effect on reticulocyte counts when fused to a fully functional wild type (WT) Fc (**Fig. S10**). This difference can be explained by the tissue specificity of the TXB2-hFc relative to the 8D3 antibody. *Ex vivo* staining of freshly isolated blood samples from untreated mice clearly showed that 8D3-hFc but not TXB2-hFc bound to reticulocytes (**Fig. S10B and C**) and consequently had no effect on circulating reticulocytes.

TXB2-hFc is a bivalent antibody that binds to recombinant mouse TfR1-ECD with 0.63 nM affinity (**Table S2**). It has been reported that high affinity, bivalent TfR1 antibodies are sorted to the lysosomes for degradation [10] and can reduce TfR1 expression in the brain endothelial cells within a day of exposure [9]. To determine whether TXB2-hFc carries a similar liability, mice were injected with TXB2-hFc at 1.875mg/kg and 18.75mg/kg (25 nmol/kg and 250 nmol/kg). TfR1 levels were quantified by Western blotting after normalization to actin and showed no observable effect on TfR1 expression 18 hr post injection even at the highest concentration (**Figs. 4 C and D**). Importantly, no adverse responses were observed in mice even with the highest dose of TXB2-hFc at 18.75mg/kg (250 nmol/kg).

### Pharmacokinetic profile

The PK properties of TXB2-hFc were further evaluated in time-course and dose-response studies. The PK profile obtained by measuring brain and plasma levels in whole brain extracts after cardiac perfusion was identical to that observed in brain fractionation studies. After a single IV dose of 1.875 mg/kg (25 nmol/kg), the brain concentration of TXB2-hFc increased exponentially over the first 18 hr and then slowly declined over the 144 hr observation period, whereas the slight increase in the concentration of control VNAR-hFc was detected a few hr after injection and remained low over the same period (**Fig. 5A**). Percentage injected dose (%ID) of TXB2-hFc in the brain at 144 hr was calculated at 3% (%ID = brain AUC/plasma AUC x 100%). Although plasma levels of the two fusion proteins diverged in the early redistribution phase, the levels were similar during the elimination phase from 18 to 144 hrs (**Fig. 5B**). In separate PK studies at 0.2 to 5 mg/kg (not shown), the terminal half-life of TXB2-hFc in plasma was calculated to be 6.5-7 days, which is expected for both human and mouse IgG in mice [25].

**Figure 5.**
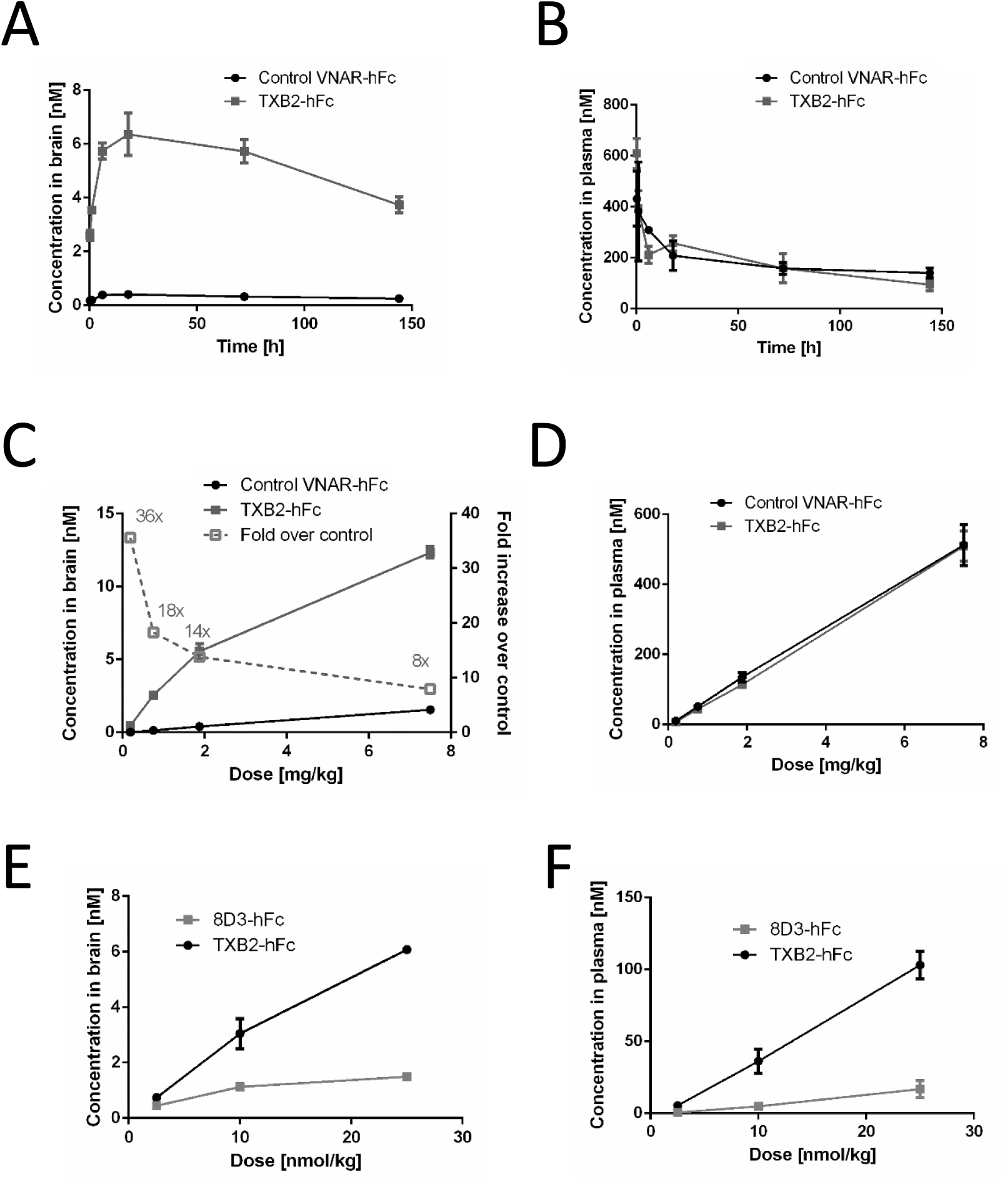
Transport kinetics of TXB2-hFc in brain after IV injection. The time course experiment with concentrations in brain (**A**) and plasma (**B**) of TXB2-hFc or control VNAR-hFc after single IV injection of 1.875 mg/kg (25 nmol/kg). Dose-response curves showing brain (**C**) and plasma (**D**) concentrations 18 hr after IV injection of TXB2-hFc or control VNAR-hFc with the fold over control indicated (dashed line) at each dose. Dose-response curves showing brain (**E**) and plasma **(F)** concentrations of TXB2-hFc compared to 8D3-hFc injected at molar equivalent doses. All tissue concentrations determined by an Fc-capture ELISA (mean +SD, N=4-5).

In dose-response studies, mice were injected IV with 0.1875, 0.75, 1.875 or 7.5 mg/kg (2.5, 10, 25 and 100 nmol/kg) of either TXB2-hFc or control VNAR-hFc and brain levels were measured 18 hr later. The greatest difference between TXB2-hFc and the control VNAR-hFc occurred at the lowest dose (over 35-fold) and exponentially decreased to 8-fold at the highest dose (**Fig. 5C)**. Conversely, the plasma levels for the two VNAR fusion proteins were identical for all doses at 18 hr after injection (**Fig. 5D**). In comparison to the 8D3-hFc antibody given at equimolar doses, TXB2-hFc resulted in significantly higher brain (**Fig. 5E**) and plasma concentrations (**Fig. 5F**). At the 25 nmol/kg dose there was over 4-fold more TXB2-hFc in the brain and 6-fold more in the plasma.

### Pharmacological response

TXB2-hFc VNAR was genetically fused to neurotensin (NT) peptide (TXB2-hFc-NT) (**Fig. S3A**), which regulates physiological responses such as body temperature from within the CNS and has been used previously as a measure of parenchymal delivery [26]. At the 25 nmol/kg IV dose, TXB2-hFc-NT significantly reduced body temperature by over 2 degrees at the 2 hr time point **(Fig. 6A**). Neither control NT fusions (control VNAR-hFc-NT or hFc-NT) produced a significant change at any of the tested time points. In the dose-response study **(Fig. 6B**), the minimal dose required to produce a hypothermic response at the 2 hr time point was 10 nmol/kg (0.75 mg/kg). Measurement of the brain and plasma concentrations of the NT fusions at the 2 and 24 hr time points showed that TXB2-hFc-NT was 10-fold higher than the control NT fusions **(Fig. 6C**) and remained constant, despite the reduction of plasma levels over the same time period **(Fig. 6D**). The recovery from hypothermia within 6 hr (**Fig. 6A**) was consistent with the development of receptor desensitization and proteolytic cleavage of the NT peptide [27].

**Figure 6.**
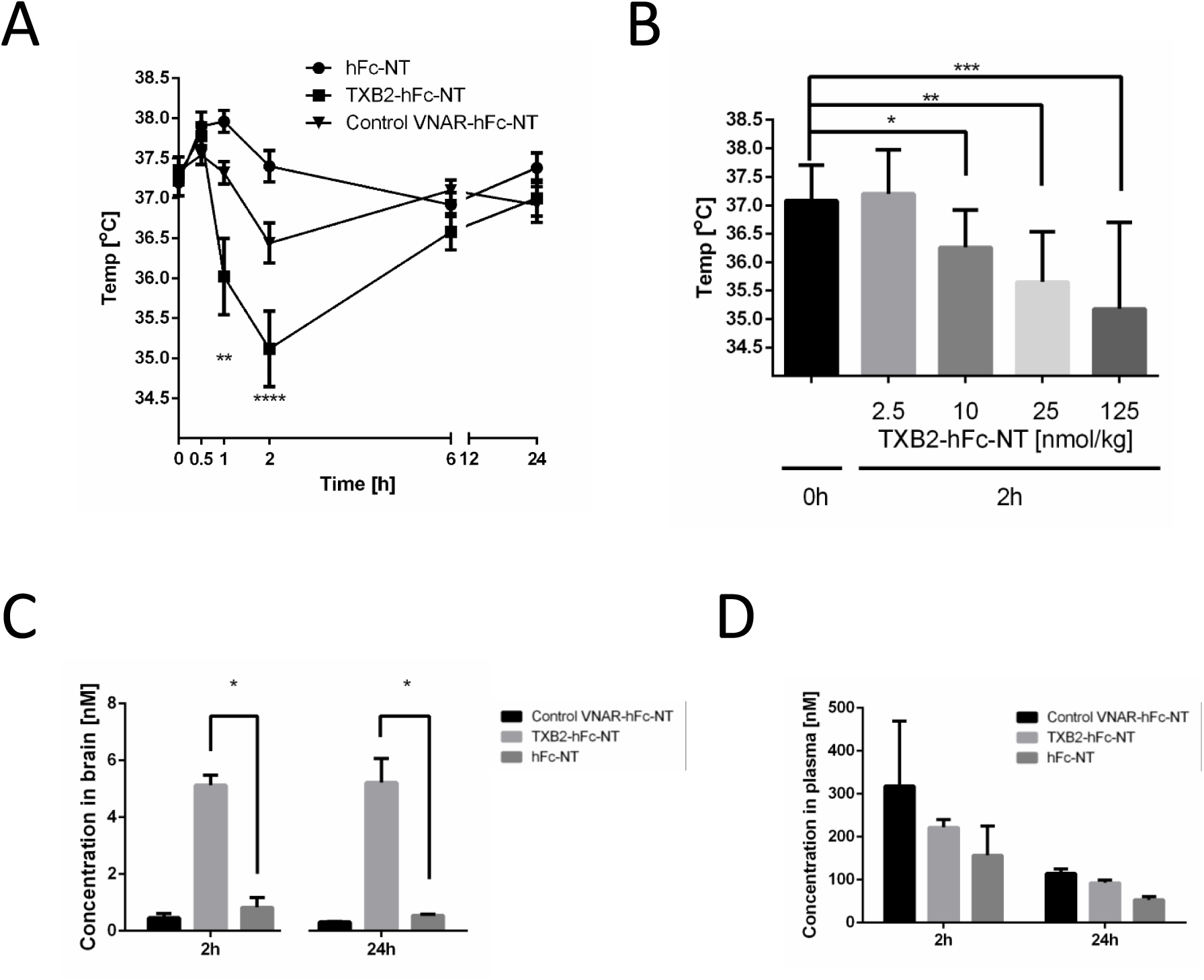
Body temperature decrease after TXB2 delivery of neurotensin (NT) to the brain. The NT peptide was genetically fused to the C-terminus of tested constructs. (**A**) Mice were injected IV with VNAR-hFc-NT fusions or hFc-NT control at 25 nmol/kg and body temperature measured over 24-hr period (mean ±SD, N=5). Significance at **-p<0.01 and ****-p<0.0001 by two-way ANOVA with Dunnett’s comparison test to body temperature measured at the beginning of the experiment for each group. (**B**) Dose-dependent hypothermia induced by TXB2-hFc-NT measured at 2 hr (mean ±SD, N=5). Significance at *-p<0.05; **-p<0.01; ***-p<0.001 determined by the Mann-Whitney test. Brain (**C**) and plasma **(D)** concentrations of the NT fusion constructs were measured by ELISA after 2 hr (N=3) and 24 hr (N=5). * Indicates significant (p< 0.01) by Student’s t-test.

## DISCUSSION

Previous efforts to identify a brain delivery vehicle using *in vivo* phage display technology attempted to target the cerebral vasculature [28-31], the CSF barrier [32, 33] or the olfactory tract [34] yet the transport mechanisms for these peptides and antibodies remain largely unknown. We combined an *in vitro* and *in vivo* selection process to isolate cross-species reactive VNARs with TfR1 specificity that are transported through the BBB into the brain parenchyma. This was achieved by first selecting VNAR phage libraries on recombinant human TfR1-ECD, injecting the pre-selected pool into the bloodstream of mice and then isolating phage clones from the brain parenchyma after capillary depletion. The change in copy number of individual VNAR sequences was tracked by NGS through the multiple rounds of the *in vitro*/*in vivo* selection process.

As anticipated, this two-tiered selection process amplified VNARs that bound both human and mouse TfR1. The most abundant unique sequences at the end of the *in vivo* selection were formatted as bivalent hFc fusion proteins (MW of 75 kDa) to increase plasma half-life and mice were treated at a low, therapeutically relevant dose (1.875 mg/kg) to identify the most potent VNARs. TXB2-hFc showed robust brain uptake, indicating that the selection process identified a BBB penetrating VNAR utilizing the TfR1 transport mechanism. Subsequent experiments confirmed this finding and further defined the biochemical and functional properties of TXB2-hFc and its potential for rapid, efficient and widespread delivery of biologics to the CNS.

TXB2-hFc bound to soluble ECDs of TfR1 from rat and cynomolgus monkey in addition to human and mouse with affinities similar to that of the reference antibodies OKT9, 8D3 and OX26 against human, mouse and rat TfR1, respectively. TXB2-hFc also recognized the receptor expressed by cells that were transiently transfected with the gene encoding either full-length mouse, human, rat or cynomolgus monkey TfR1. Importantly, TXB2-hFc did not compete with Tf or ferritin for TfR1-ECD binding in biochemical assays. These results suggest that TXB2 binds to a surface accessible epitope on TfR1 that is highly conserved and distant from the Tf/ferritin binding interface. While species cross-reactive antibodies to the surface regions of TfR1 are rare, the finding that species cross-reactive VNARs are common could be related to their ability to access buried epitopes [35].

The relationship between TfR1-ECD binding affinity and brain penetration for TXB2 stands in contrast to what has been found with various reference antibodies to the receptor. High affinity binding of antibodies to TfR1 appears to impede transcytosis by lysosomal targeting or preventing receptor dissociation consequently trapping the antibody inside the capillary endothelium [8, 10,36-38]. It was further shown that monovalent formatting or lowering the affinity of TfR1 antibodies could boost brain transport *in vivo* [8, 11-13]. Conversely, a series of biochemical, histological, biodistribution, pharmacokinetic and pharmacodynamic studies showed that neither high-affinity binding (0.6 nM KD) nor a bivalent configuration impeded BBB transport of TXB2-hFc via TfR1.

### Brain distribution

The brain distribution of TXB2-hFc fusion after IV injection was first examined by biochemical fractionation studies. The results indicate that TXB2-hFc accumulated in the brain parenchyma like TXB2 expressed on the phage surface, thereby validating the *in vivo* phage selection approach. Furthermore, tracking the distribution of TXB2-hFc after a single dose over time indicates BBB penetration occurs through a rapid, saturable transport mechanism. The time-course study showed that TXB2-hFc levels in the capillary fraction measured 15 min after injection remained constant for at least 72 hr. By contrast, levels in the parenchyma fraction rapidly rose to exceed the capillary levels by 18 hr post-injection and thereafter the two fractions maintained a constant equilibrium throughout the 6-day study.

IHC studies confirmed the parenchymal distribution of TXB2-hFc after IV administration. In addition to the prominent staining of entire microcapillary endothelium, there was also a diffuse widespread staining of the brain parenchyma and specific localization within multiple neuronal populations. The neuronal distribution pattern mirrored TfR1 expression with the most intense TXB2-hFc staining seen in the brain stem and cerebellum, which likely reflects uptake in neurons with highest TfR1 expression [23, 24]. Double immunofluorescent staining confirmed the co-localization of TXB2-hFc with TfR1-positive neurons and a striking absence of TXB2-hFc immunoreactivity in astrocytes, microglia and oligodendrocytes, which is in accordance with observations on absence of TfR1 on these cell types [19, 23]. Despite its high affinity and bivalent format, TXB2-hFc shows more robust staining than previously reported for TfR1 antibodies to date [8, 11], and the slow clearance of TXB2-hFc sequestered in neurons most likely accounts for its long half-life in the brain.

The specificity of TXB2-hFc in brain was confirmed by *ex vivo* IHC staining of paraffin sections prepared from untreated mice. The staining pattern for TXB2-hFc in various brain regions including the cerebral cortex, hippocampus, anterior thalamus, striatum and cerebellum was nearly identical to the 8D3 reference antibody. However, the control VNAR-hFc, which binds the soluble ECD and full-length receptor in transfected cells, failed to react with TfR1 on brain sections just like an irrelevant VNAR, which explains why it cannot penetrate the BBB.

### Biodistribution

Access of antibodies to peripheral tissues through capillary leakage and diffusion can affect their biodistribution, pharmacokinetics and off-target effects. TfR1 is mainly expressed by cells in the body with high iron demand including lymphoid cells in the spleen, hepatocytes in the liver, myocytes in the heart and skeletal muscle, renal tubules in the kidney and epithelial cells and macrophages in the lung [1]. After IV injection, there was a dramatic accumulation of TXB2-hFc in the brain whereas the levels found in all peripheral organs sampled except for the lung were similar to the VNAR controls. In comparison, the 8D3 and RI7217 TFR1 antibodies were reported to have a different biodistribution pattern in mice. RI7217 accumulates in the brain and heart, but is not taken up by the liver or kidney, whereas 8D3 antibody accumulates in liver and kidney in addition to the brain [6, 7]. Another monoclonal antibody to TfR1 called TSP-A18 preferentially accumulated in spleen and bone but not in other mouse organs [39]. Thus, differences in tissue selectivity between various TfR1 antibodies indicates that the presentation and accessibility of TfR1 epitopes *in vivo* is context dependent.

TXB2-hFc appears highly selective for brain relative to other TfR1 antibodies characterized to date. An independent study using radiolabeled TXB2-hFc corroborated the selective distribution to brain after intravenous administration with no increase over control in any other organ except lung [52]. We also confirmed that unlike the 8D3 antibody, TXB2 does not recognize TfR1 expressed on mouse reticulocytes. These studies indicated that the TfR1 binding epitope for TXB2-hFc was inaccessible in most peripheral organs, and thus less likely to target a cargo to peripheral tissues. In addition, TXB2-hFc is less likely to undergo tissue-mediated clearance, which may account for the long plasma half-life relative to other TfR1 antibodies. Additional studies are needed to confirm and further characterize the cellular distribution and consequence of increased levels of TXB2-hFc in lung.

### Safety profile

Historically TfR1 antibodies were generated with cytotoxic activity to target proliferating tumor cells that overexpress the receptor [40]. To avoid potential toxicity associated with a BBB shuttle, we and others have selected antibodies that do not block the binding of endogenous Tf to its receptor, which could lead to iron deprivation in the CNS [19]. TfR1 antibodies indirectly prevent iron uptake by interfering with TfR1 recycling and trafficking, which has resulted in lysosomal degradation of the receptor in the capillary endothelium [9, 10]. Reducing receptor binding affinity or monovalent formatting of these antibodies restored intracellular trafficking and maintained TfR1 levels in the CNS. Furthermore, exposure to TfR1 antibodies with full effector function caused severe anemia in mice due to the hemolysis [4] and immune-mediated CNS toxicity in addition to anemia in primates [5]. However, it was previously shown that direct antibody-medicated cytotoxicity was related to effector function, which can be attenuated by engineering the antibody Fc region [4, 18, 41]. TXB2-hFc was evaluated in light of these potential safety liabilities, and we found that it did not block Tf or ferritin binding or reduce TfR1 levels in the CNS. Unlike the 8D3-hFc antibody, TXB2-hFc did not bind reticulocytes *ex vivo* and consequently did not deplete them from the circulation, with or without Fc effector mutations. This finding is in stark contrast to the robust loss of circulating reticulocytes seen with monoclonal antibodies to TfR1 administered at similar doses, which is only partially reversed by the elimination of effector function due to residual activation of the complement pathway [4, 42].

### Pharmacokinetics

The PK profile of TXB2-hFc is an important distinguishing feature relative to other TfR1 antibodies used as BBB shuttles. Robust brain levels were achieved over an extended period after a single low dose (1.875 mg/kg). The long half-life in the brain can be explained by continuous uptake from plasma via a receptor-mediated transport mechanism and sequestration in TfR1-positive neurons. It should also be noted that measurements of TXB2-hFc in whole brain extracts underestimates the ratio relative to controls since the control VNAR-hFc was not found in the brain capillaries or parenchymal by IHC. Additionally, there was no evidence of a specific clearance mechanism as the plasma elimination rate of TXB2-hFc was as expected for IgG. By contrast, other high affinity TfR1 antibodies have a short plasma half-life indicative of a target-mediated clearance mechanism; mutations that reduce antibody binding affinity extend the plasma half-life although not to the level of untargeted IgG [4, 6, 8, 11, 13]. We likewise confirmed the significantly lower brain uptake and short plasma half-life of the 8D3-hFc antibody in head-to-head experiments with TXB2-hFc.

While the IHC studies provide strong evidence that TXB2-hFc is principally transported through the BBB pathway, it is possible that BCSFB transport provides a minor contribution to the levels measured in whole brain extracts in our PK studies. Epithelial cells of the choroid plexus express TfR1 in addition to many other transporters found in the BBB and provide active endocytic pathways into the CSF [43-45]. Additionally, PK studies using microdialysis revealed a nonselective pathway for peripherally administered antibodies into the brain via BCSFB into the ventricles [46]. Therefore, the relative contribution of both BCSFB and BBB pathways to brain delivery via BBB shuttles warrants consideration.

### CNS activity

Using neurotensin (NT) as a cargo has provided a convenient physiological readout to monitor the efficacy of brain delivery. Endogenous NT is produced by neurons and glial cells within the brain where it functions as a neurotransmitter to regulate a wide array of physiological processes, including body temperature, blood pressure and nociception from within the CNS [47]. In mice, intracisternal delivery of 10-30 µg NT maximally reduced body temperature within 30 min, which returned to near normal 2 hr later [48]. IV delivery in the rat of NT fused to Angiopep-2 (a peptide targeting LDL receptor–related protein-1, LRP1) at a dose of 1.3 µmol/kg produced a more potent and extended response than CSF delivery due to the prolonged plasma half-life of the NT fusion protein [26]. By comparison, the present study showed that IV delivery of NT fused to TXB2-hFc at 25 nmol/kg in mice produced a similar reduction in body temperature that peaked 2 hr after administration and returned to normal by 6 hr. The onset of the response is consistent with the observed transit time for TXB2 across the BBB in brain fractionation studies while the duration of hypothermia induced by NT is limited by both receptor desensitization [49] and proteolytic cleavage [27]. Interestingly, the potency of TXB2-hFc-NT was 50-fold greater than Angiopep-2-NT in lowering body temperature, which may reflect the ability of the TXB2-hFc shuttle to deliver the cargo across the BBB and directly target TfR1-expressing neurons. The use of mild to moderate pharmacologically induced hypothermia has shown potential in traumatic brain injury and stroke in various experimental models and in clinical trials [50, 51]. Therefore, the use of targeted delivery of neurotensin to the brain by TXB2 via IV administration might provide a significant improvement to induction of hypothermia.

In summary, the properties of VNAR TXB2 clearly differ in several important aspects from those of monoclonal antibodies to TfR1 used as BBB shuttles. In contrast to high-affinity TfR1 antibodies including 8D3, OX26, and anti-TfR^A^[8], high affinity binding of TXB2 did not impede transport across the BBB. Subsequent to capillary binding, TXB2-hFc was widely distributed across the brain and was internalized by TfR1-expressing neuronal populations. Moreover, the pharmacokinetic profile of TXB2-hFc was distinguished by a more selective biodistribution, prolonged brain exposure, and the lack of target-mediated clearance. Consequently, the minimal effective doses of TXB2-hFc fused to NT was as low as 1-2 mg/kg (12.5-25 nmol/kg), indicating that a high-affinity shuttle is capable of efficient BBB transport. While TXB2 clearly differentiate from other TfR1 antibody shuttles as a result of the functional *in vivo* selection, further mechanistic studies are needed to determine whether its favorable profile derives from the inherent properties of a VNAR, the particular TfR1 epitope to which it binds or a combination of these factors.

## ABBREVIATIONS

(TfR1): Transferrin Receptor 1
(BBB): blood-brain barrier
(Tf): Transferrin
(CNS): central nervous system
(VNAR): Variable domain of New Antigen Receptors
(CDR3): complementarity-determining region 3
(RT): room temperature
(SEC): size exclusion chromatography
(HSA): human serum albumin
(NT): Neurotensin
(IHC): Immunohistochemistry
(NGS): next generation sequencing
(PK): pharmacokinetic
(BCSFB): blood-CSF barrier
(%ID): Percentage injected dose
(AUC): Area under the curve
(AEF): attenuated effector function
(SDS-PAGE): sodium dodecyl sulfate polyacrylamide gel electrophoresis
(WB): Western blot
(IV): intravenous
(ECD): ectodomain

## ACKNOWLEDGMENTS

The authors wish to thank Martin Flajnik for valuable insight into the VNAR single domain and Hanne Krone Nielsen and Merete Fredsgaard for excellent technical assistance.

## Supplementary Material

**Table S1.** The relative abundance of top 29 clones after selection by NGS. VNAR sequences are grouped in families with common CDR3s and the 29 most abundant families are shown. Relative abundance was calculated as per million sequences at each stage of the selection.

|  | OsX-3/4 | in vitro R1 | in vitro R2 | in vivo R1 | in vivo R2 | in vivo R3 |
| --- | --- | --- | --- | --- | --- | --- |
| A | 0 | 0 | 808 | 1163 | 1329 | 996 |
| B | 0 | 0 | 306 | 533 | 686 | 709 |
| C | 0 | 1 | 210 | 360 | 506 | 616 |
| D | 0 | 0 | 122 | 184 | 235 | 167 |
| E | 0 | 0 | 93 | 163 | 166 | 205 |
| F | 0 | 0 | 99 | 158 | 214 | 177 |
| G | 0 | 0 | 100 | 154 | 174 | 200 |
| H | 0 | 0 | 50 | 76 | 108 | 88 |
| I | 0 | 0 | 38 | 65 | 79 | 89 |
| J | 0 | 0 | 50 | 58 | 93 | 86 |
| K | 0 | 0 | 38 | 52 | 69 | 91 |
| L | 0 | 0 | 45 | 50 | 98 | 79 |
| M | 0 | 0 | 16 | 48 | 49 | 53 |
| N | 0 | 0 | 22 | 45 | 71 | 55 |
| O | 0 | 0 | 34 | 41 | 44 | 31 |
| P | 0 | 0 | 23 | 35 | 38 | 15 |
| Q | 0 | 0 | 22 | 35 | 36 | 19 |
| R | 0 | 0 | 13 | 33 | 67 | 71 |
| S | 0 | 0 | 16 | 32 | 45 | 46 |
| T | 0 | 0 | 10 | 30 | 45 | 62 |
| U | 0 | 0 | 9 | 30 | 42 | 64 |
| V | 0 | 0 | 10 | 28 | 36 | 24 |
| W | 0 | 0 | 23 | 26 | 40 | 21 |
| X | 0 | 0 | 17 | 26 | 43 | 48 |
| Y | 0 | 0 | 16 | 22 | 23 | 9 |
| Z | 0 | 0 | 18 | 20 | 55 | 57 |
| AA | 0 | 0 | 10 | 20 | 44 | 33 |
| AB | 0 | 0 | 1 | 13 | 37 | 50 |
| AC | 0 | 5 | 11 | 19 | 26 | 34 |

**Table S2.** Binding kinetics to recombinant human, mouse, rat and cynomolgus monkey TfR1-ECD (hTfR1, mTfR1, rTfR1 and cTfR1). All TfR1 constructs were expressed and purified in-house. SPR assays were performed using single cycle kinetics on an anti-His capture-chip coated with His-tagged TfR1 at 10 µg/ml. The association rate (ka), dissociation rate (kd) and binding affinity (KD) for TXB2-hFc and control VNAR-hFc were calculated. Holo-hTf specific for human TfR1 as well as control antibodies were included with 8D3, OX26, and OKT9, specific to mouse, rat and human, respectively.

| Analyte | hTfR1 |  |  | mTfR1 |  |  | rTfR1 |  |  | cTfR1 |  |  |
| --- | --- | --- | --- | --- | --- | --- | --- | --- | --- | --- | --- | --- |
| | $k_a$ (1/Ms) | $k_d$ (1/s) | KD (M) | $k_a$ (1/Ms) | $k_d$ (1/s) | KD (M) | $k_a$ (1/Ms) | $k_d$ (1/s) | KD (M) | $k_a$ (1/Ms) | $k_d$ (1/s) | KD (M) |
| control VNAR-hFc | 5.1E+4 | 8.6E-5 | 1.7E-9 | 4.7E+4 | 7.3E-5 | 1.5E-9 | ND | ND | ND | 4.9E+6 | 1.5E-3 | 3.1E-10 |
| TXB2-hFc | 2.2E+5 | 8.6E-5 | 3.9E-10 | 1.6E+5 | 1.0E-4 | 6.3E-10 | 1.0E+5 | 1.2E-4 | 1.2E-9 | 2.6E+5 | 7.4E-5 | 2.8E-10 |
| Holo hTf | 6.8E+4 | 4.6E-4 | 6.8E-9 | ND | ND | ND | ND | ND | ND | ND | ND | ND |
| 8D3-hFc | ND | ND | ND | 4.1E+5 | 7.3E-5 | 1.8E-10 | ND | ND | ND | ND | ND | ND |
| OKT9 | 1.5E+5 | 1.9E-5 | 1.3E-10 | ND | ND | ND | ND | ND | ND | ND | ND | ND |
| OX26 | ND | ND | ND | ND | ND | ND | 7.7E+4 | 1.5E-4 | 2.0E-9 | ND | ND | ND |

**Figure S1.**
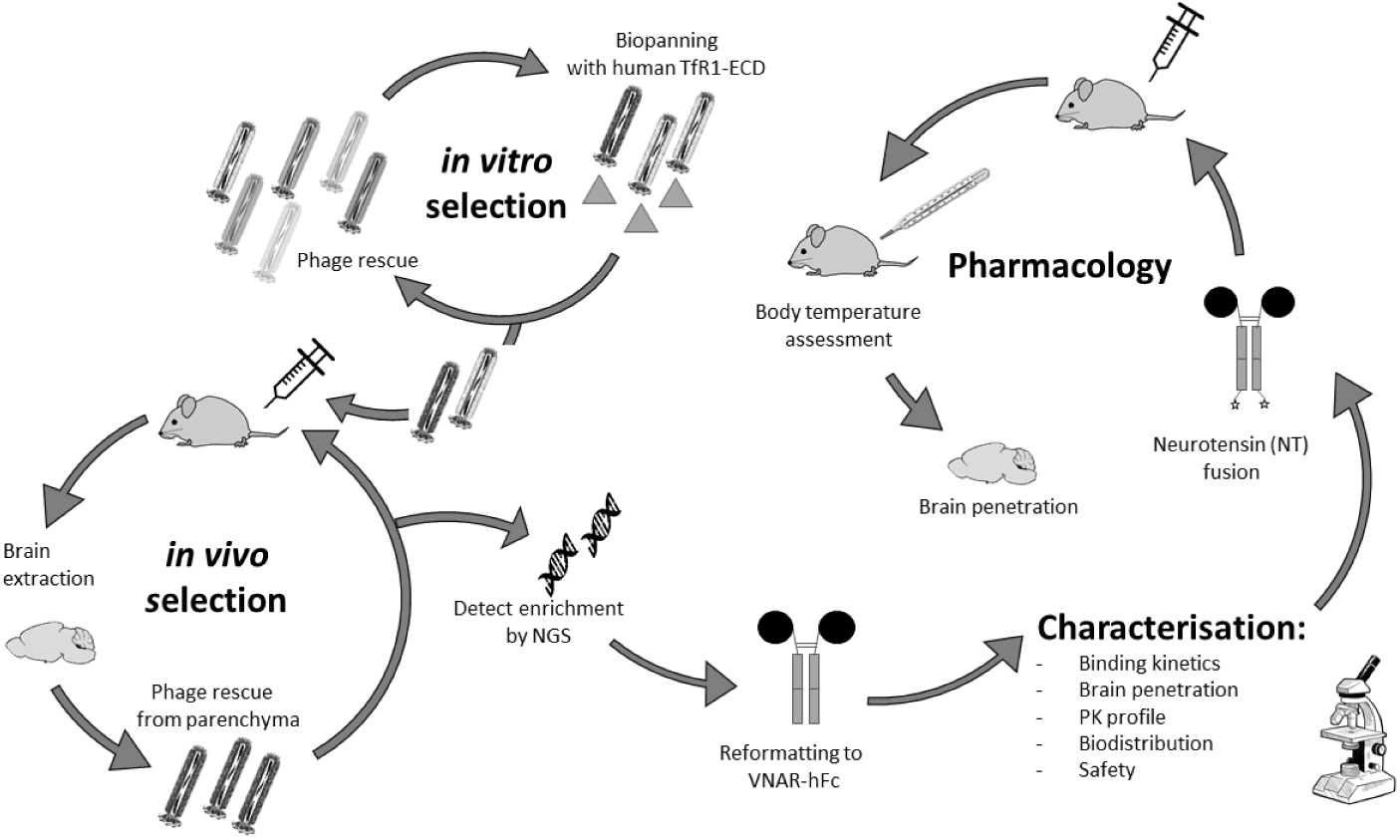
Overview of the VNAR phage library selection and screening process for the identification of the TXB2 BBB shuttle. Steps include *in vitro* and *in vivo* selections, reformatting as Fc fusion proteins and lead characterization followed by pharmacological assessment of payload delivery to the brain.

**Figure S2.**
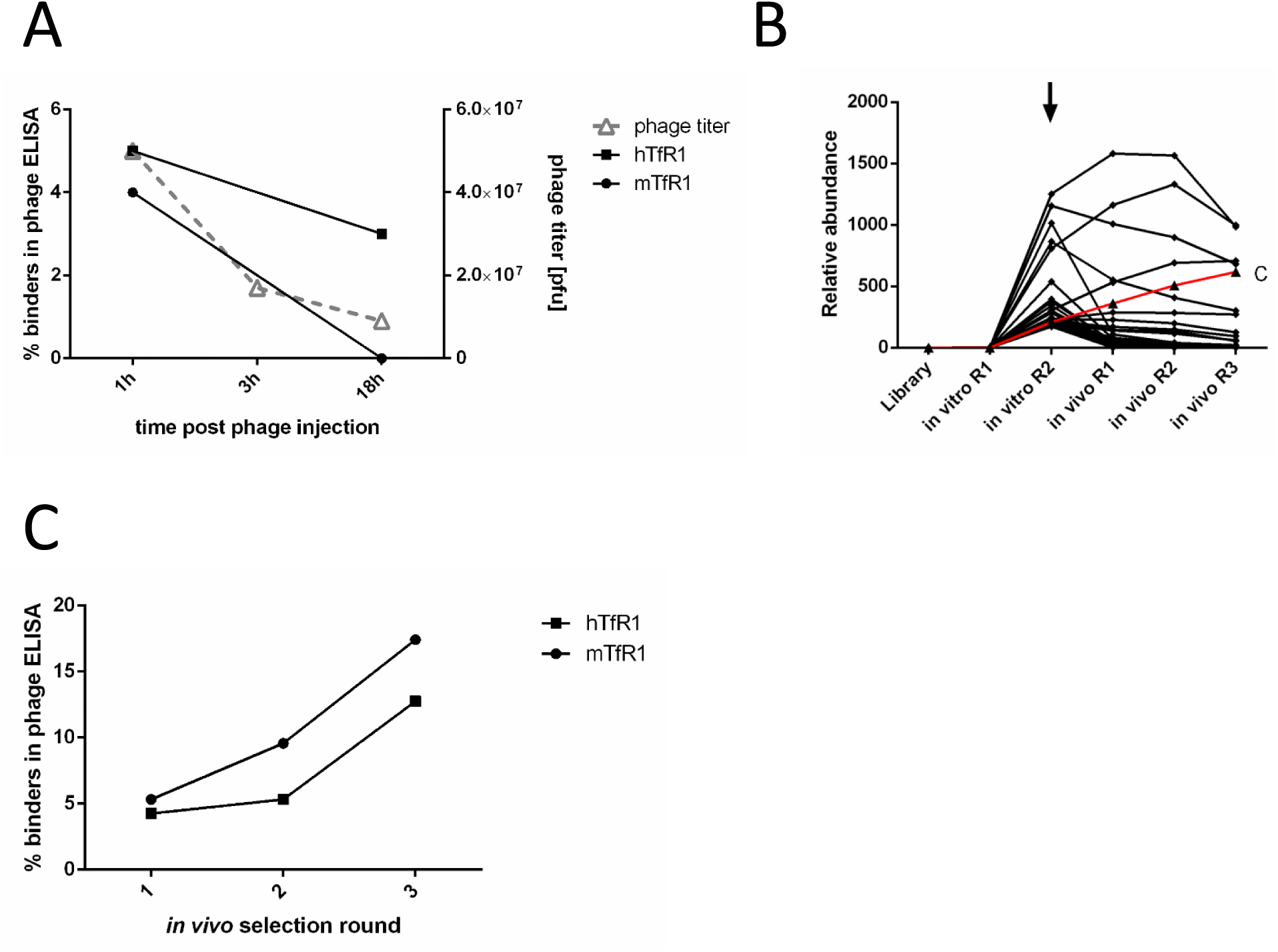
In vitro/in vivo selection of brain penetrant VNARs. (**A**) Optimization of in vivo selection method for TfR1-dependent brain penetrants. Preselected phage library was injected to animals and the brains were extracted 1, 3 or 18 hr post injection following cardiac perfusion. The phage output titer was calculated using plaque-forming unit (pfu) in parenchymal fraction only. Approximately 100 clones were randomly picked from 1- and 18-hr time-points and binding to human (■) and mouse (•) TfR1-ECD was assessed by phage ELISA. (**B**) The 20 most abundant sequences at the 2nd in vitro round of selection (marked by arrow) were tracked for their abundance in the further rounds of *in vivo* selection. (**C**) Individual clones were randomly picked and tested in phage ELISA for binding to human and mouse TfR1-ECD (hTfR1 and mTfR1). Approximately 200 clones were picked for each of rounds 1 and 2, and 800 clones were picked for round 3. A clone was defined as a binder if its signal exceeded four times the background signal.

**Figure S3.**
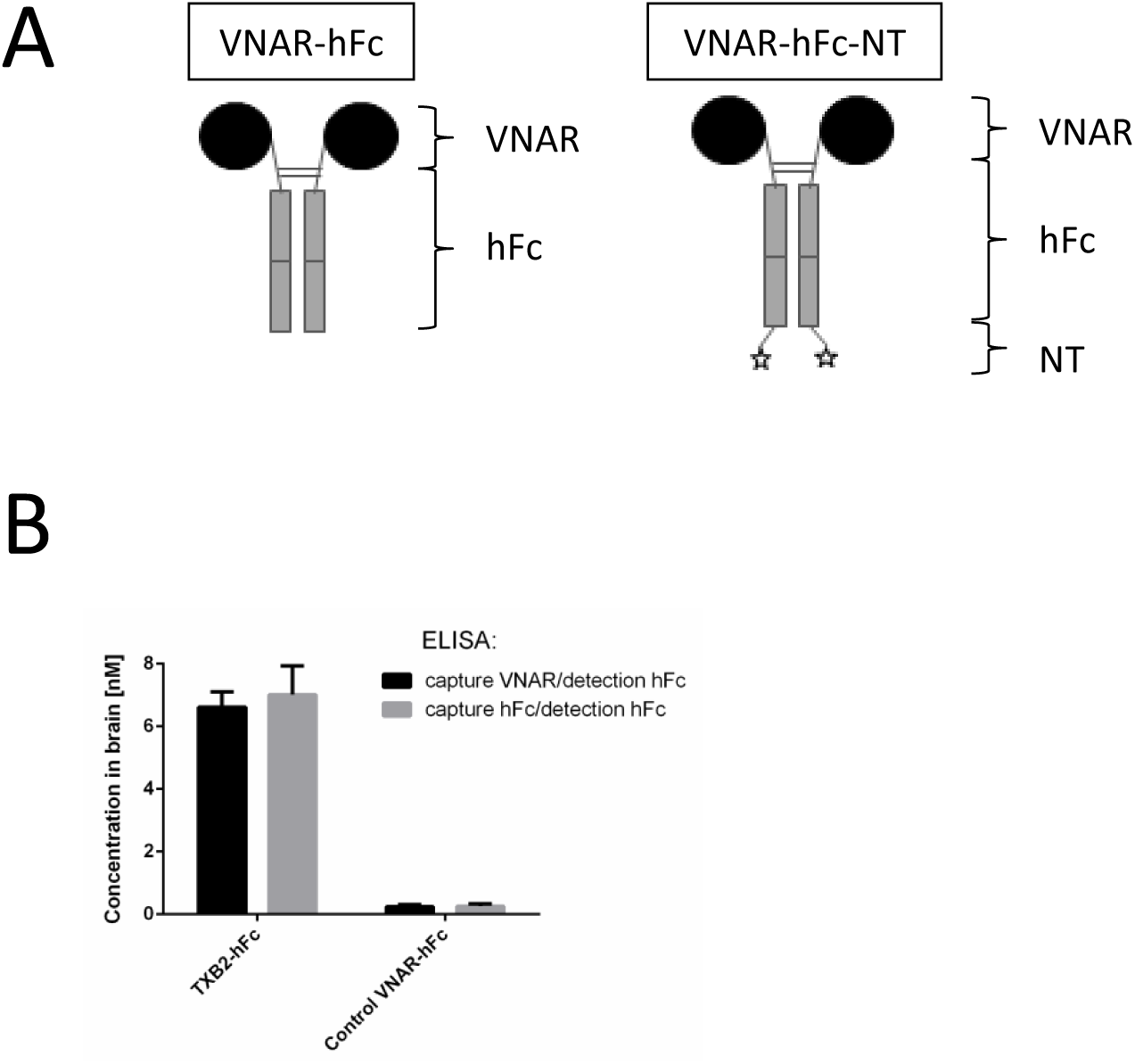
Cartoon depiction of bivalent VNAR-fusion proteins and alternate ELISA detection methods. (**A**) TXB2-hFc and TXB2-hFc-NT formats, respectively. (**B**) Fc-capture or VNAR-capture using the same Fc detection antibody shows the stability of the VNAR-Fc format in mouse brain by ELISA (mean ±SD, N=3).

**Figure S4.**
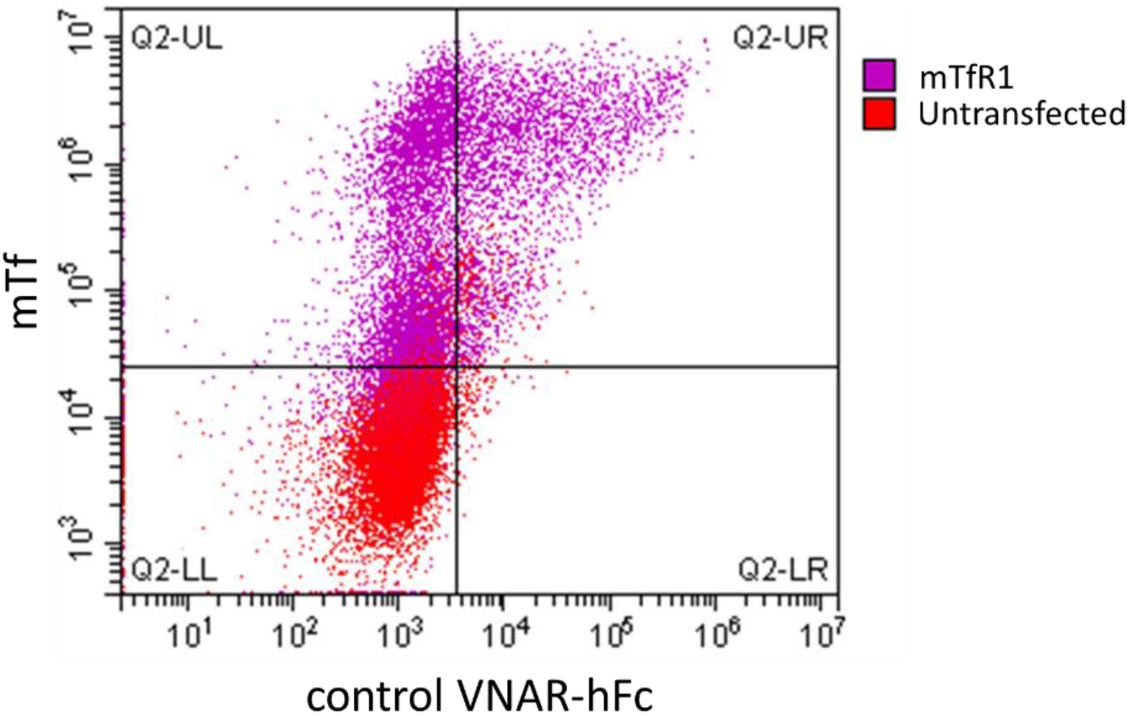
Binding of control VNAR-hFc to transiently expressed mTfR1. Live HEK293 cells transiently transfected with mTfR1 were double stained with the control VNAR-hFc and mouse transferrin (mTf) and analyzed by flow cytometry. Double-positive cells that populated Q2 after transfection showed binding of both control VNAR-hFc and mTf to the transiently expressed mouse TfR1.

**Figure S5.**
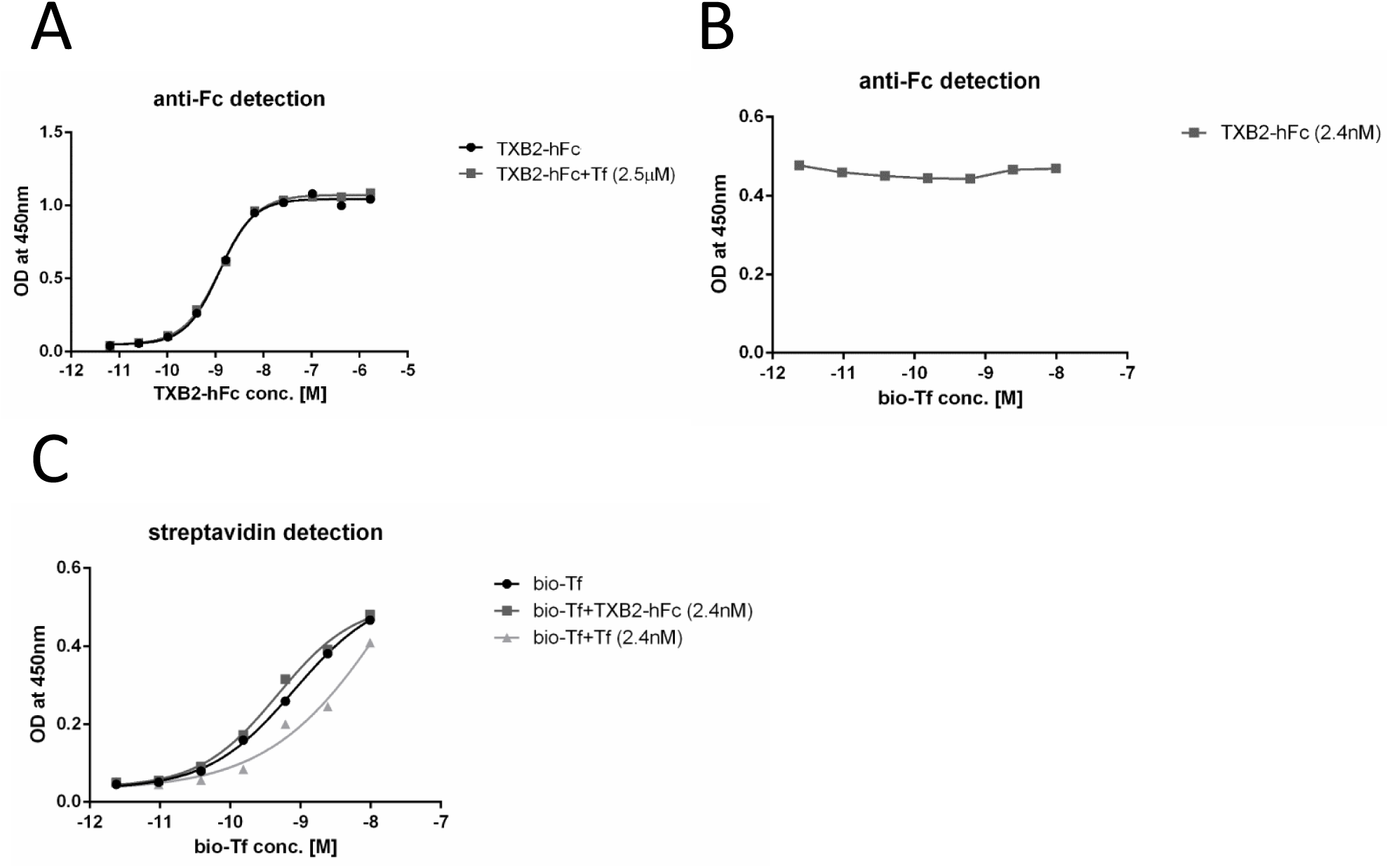
Competition analysis between TXB2-hFc and Tf for binding to rh-TfR1-ECD. ELISA plates were coated with rh-TfR1-ECD. (**A**) Binding of TXB2-hFc to rh-TfR1-ECD at concentrations ranging from pM to µM was assessed on its own and after pre-incubation with 2.5 µM of recombinant Tf. (**B** and **C**) Alternatively, plates were pre-incubated with biotinylated Tf (bio-Tf) at concentrations ranging from pM to µM. After the addition of either TXB2-hFc or unlabeled Tf (both at 2.4 nM), TXB2-hFc binding was detected with anti-human Fc antibody (**B**) and bio-Tf with streptavidin (**C**).

**Figure S6.**
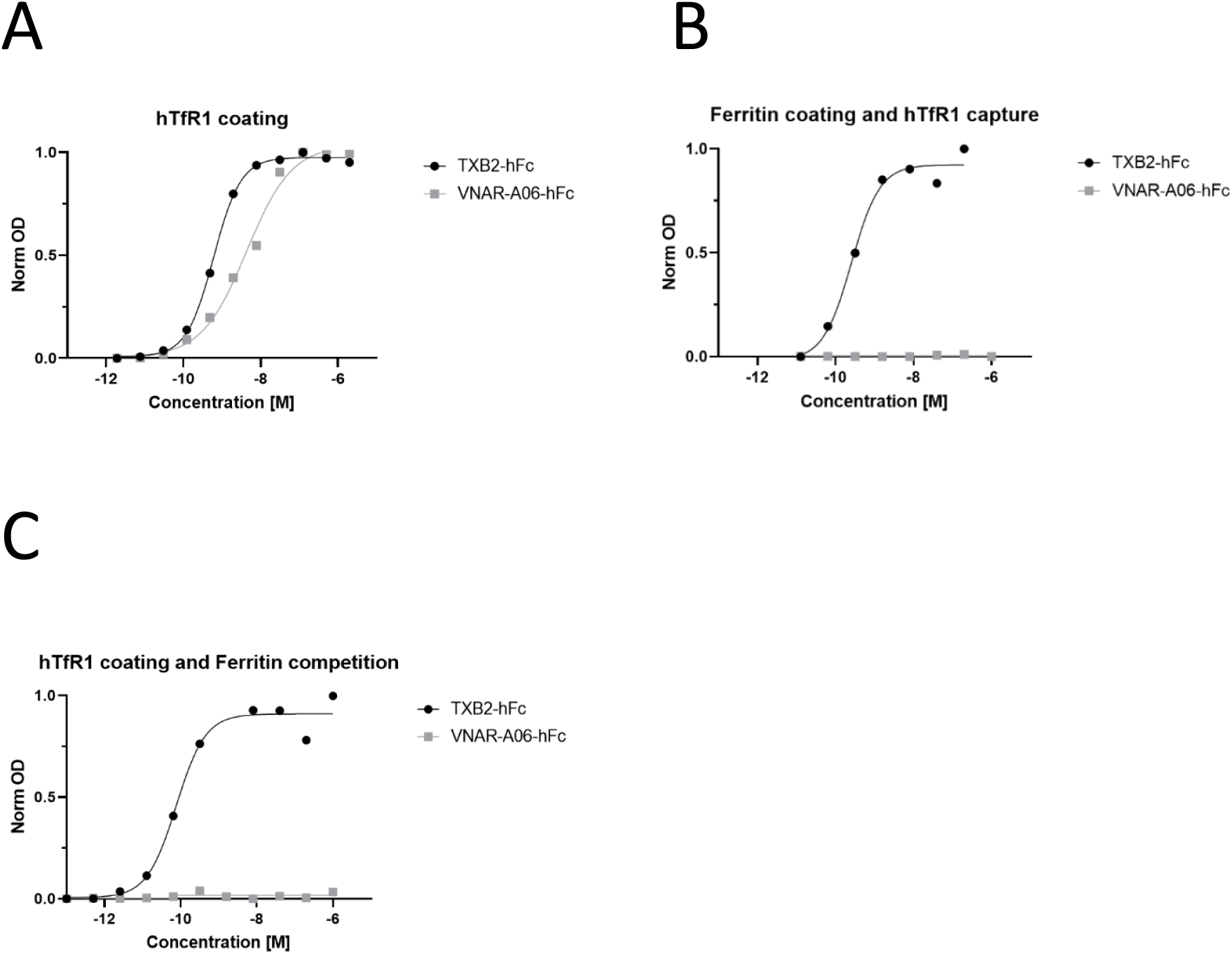
Competition analysis between TXB2-hFc and ferritin for binding to rh-TfR1-ECD. (**A**) ELISA plates were coated with rh-TfR1-ECD. Binding of TXB2-hFc and VNAR-A06-hFc (control VNAR antibody) to rh-TfR1-ECD was assessed at concentrations ranging from pM to µM. (**B**) ELISA plates were coated with ferritin, which was used to captured rh-TfR1-ECD. Binding of TXB2-hFc and VNAR-A06-hFc to rh-TfR1-ECD was then assessed at concentrations ranging from pM to µM. (**C**) ELISA plates were coated with rh-TfR1-ECD and serial dilutions of TXB2-hFc or VNAR-A06-hFc were added in the presence of excess ferritin (100 nM). Absorbance was normalized to the highest individual point and presented as normalized optical density (Norm OD).

**Figure S7.**
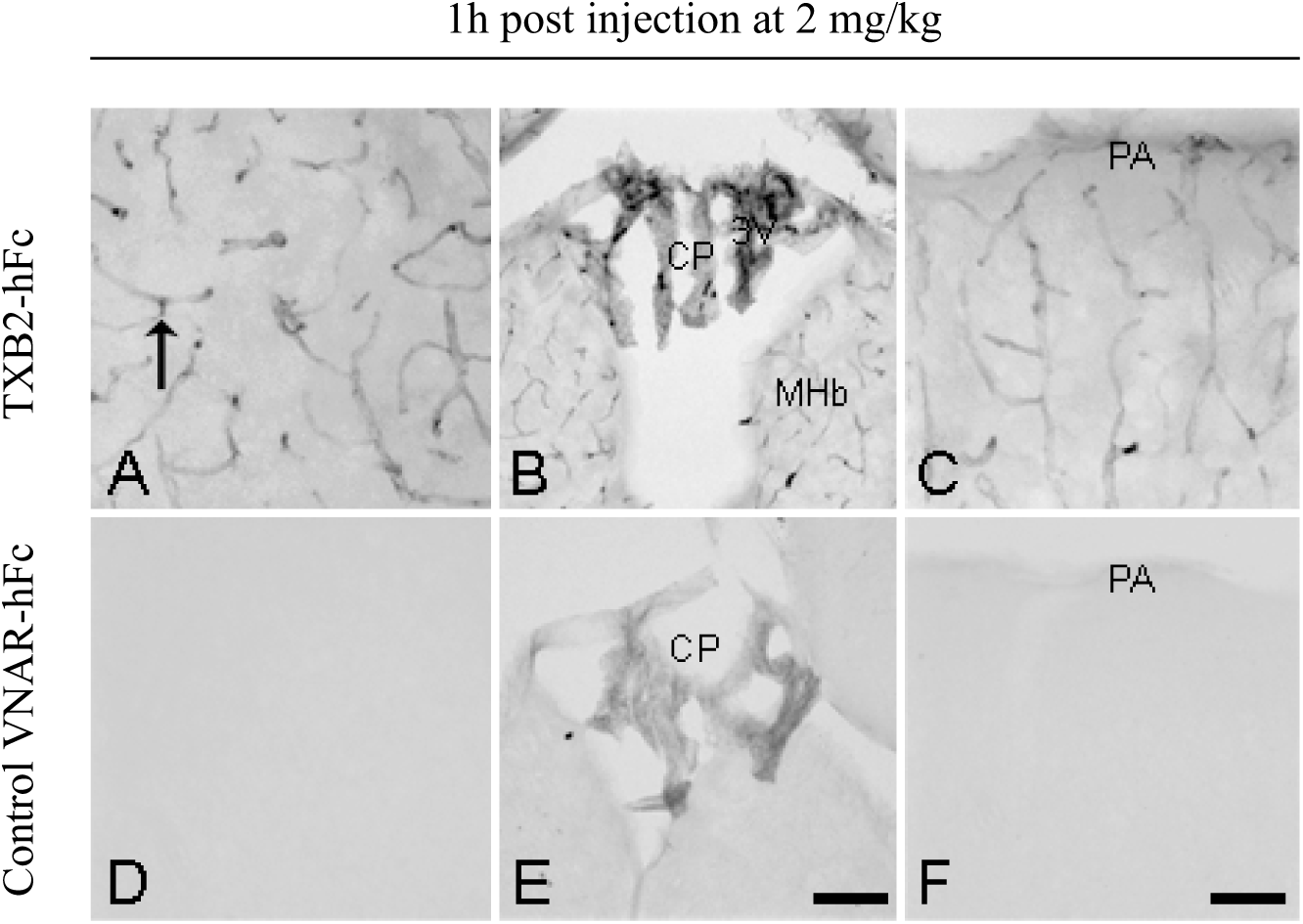
The distribution of TXB2-hFc evaluated using IHC in the brain vascular. (**A-C**) TXB2-hFc injected in a dose of 2 mg/kg, IV and allowed to circulate for 1 hr. (**A**) TXB2-hFc is present in brain capillaries (arrow) throughout the brain. (**B)** Labelling is also seen in the choroid plexus (CP), but not in the adjacent brain tissue containing the medial habenular nucleus (MHb) lining the third ventricle (3V). (**C)** In the cerebral cortex and the surface of the brain, labelling is confined to brain capillaries and is not prominent at the level of the pia-arachnoid membrane (PA). (**D-F**) Control VNAR-hFc injected at a dose of 2 mg/kg and allowed to circulate for 1 hr. (**D**) Labelling is virtually absent and does not discern any pattern of labelling in brain capillaries. (**E**) Labelling is nonetheless seen in the CP, which can be attributed to non-specific transport at the BCSFB. (**F**) Labelling is virtually absent from the PA, indicating that the amount of control VNAR-hFc transported into the brain is low. Scale bars: **A, C, D, F** 50 µm (bar shown in **F**), **B, E** 200 µm (bar shown in **E**).

**Figure S8.**
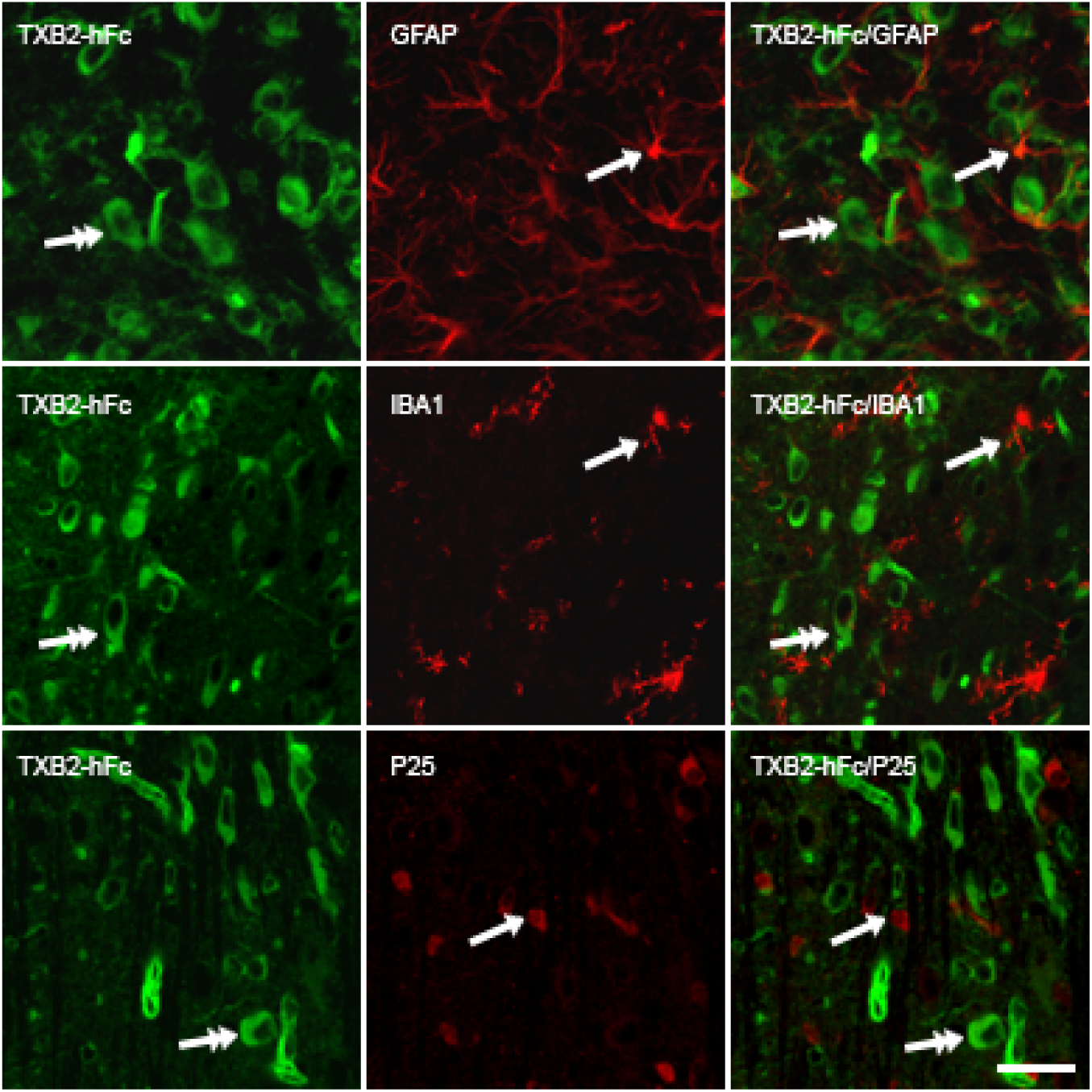
TXB2-hFc did not co-localize with glial cells in the brain after IV injection. Distribution of TXB2-hFcwas evaluated by double-labelling to simultaneously detect the major glial cell types astrocytes (upper row), microglia (middle row), and oligodendrocytes (lower row) in high magnification. Sections taken from mesencephalon (astrocytes), lateral (microglia) and medial (oligodendrocytes) portions of the pons. The co-detection reveals that the various glial markers (red, single arrow) identifying numerous glial cells do not co-localize with TXB2-hFc (green, double-headed arrows). Scale bar: 80 µm.

**Figure S9.**
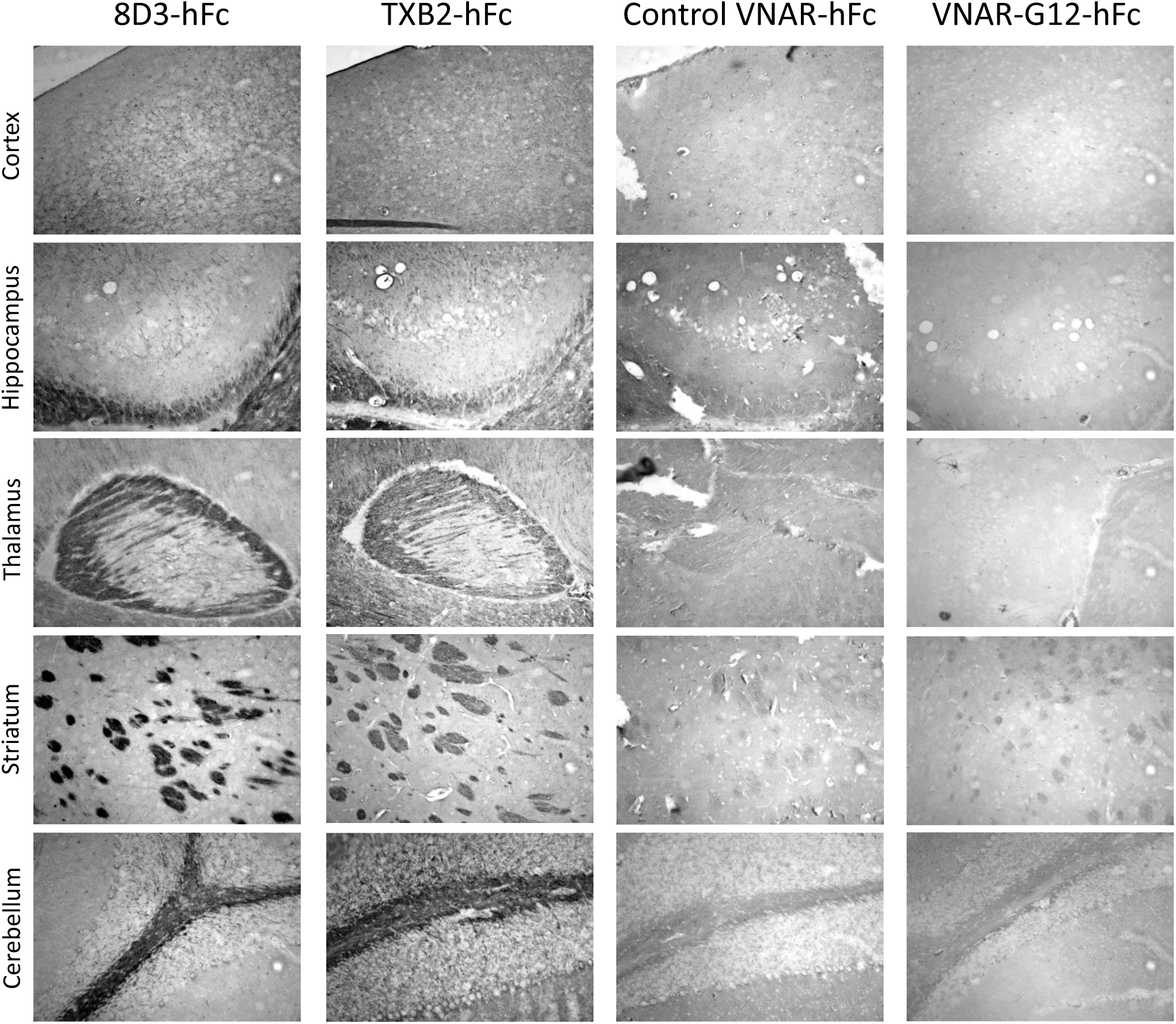
IHC *ex vivo* staining of mouse brain section after direct application of TXB2-hFc. Staining was compared to 8D3-hFc and two VNAR-hFc controls (TfR1-ECD binding control and TfR1 non-binding VNAR G12). Paraffin embedded sections from untreated mice were incubated with primary antibodies overnight at 4°C and binding was detected with biotinylated-anti-human IgG and avidin-HRP. All pictures were taken at 20X magnification with matched parameters for acquisition.

**Figure S10.**
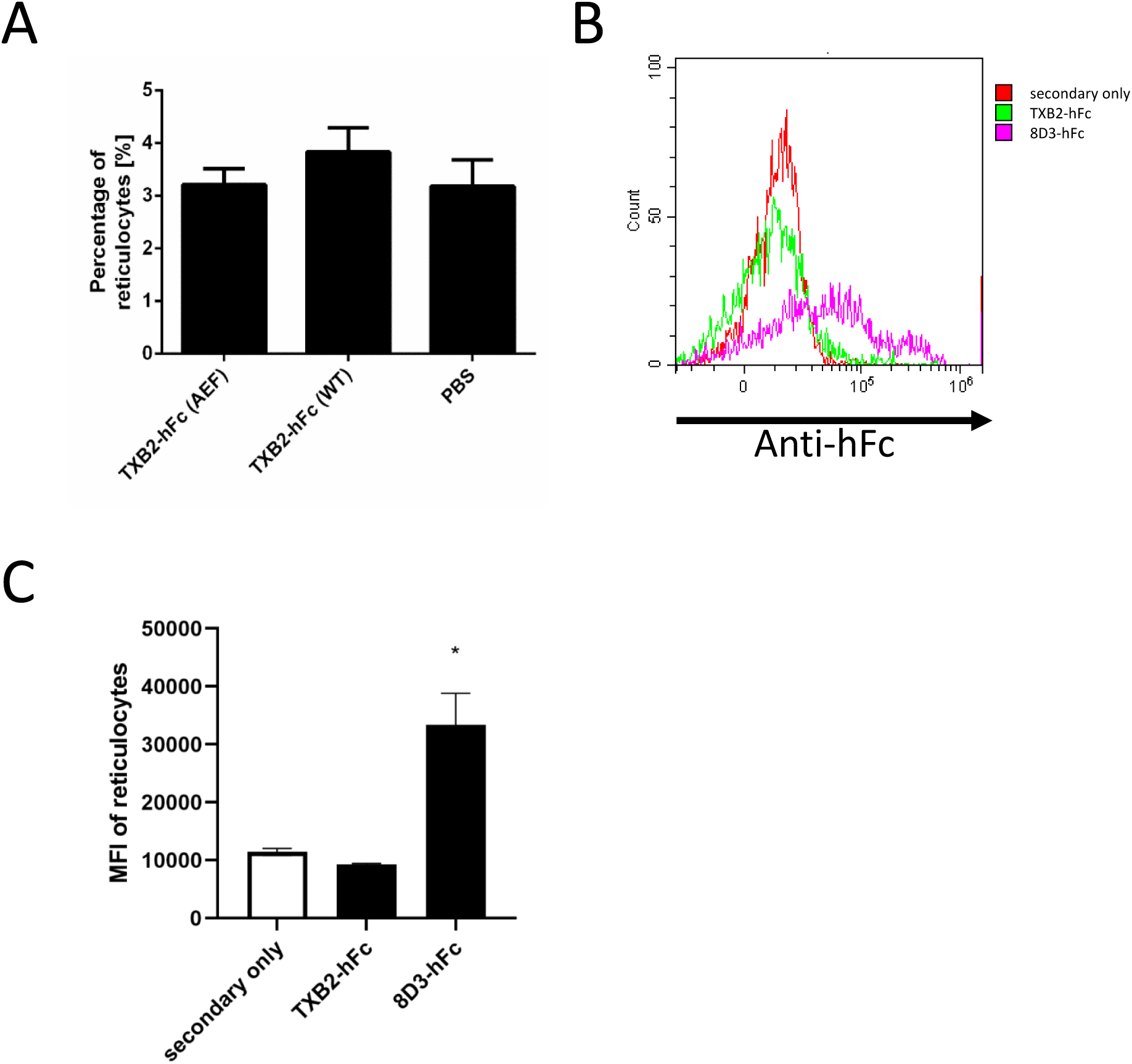
TXB2-hFc does not deplete reticulocytes *in vivo* or bind to reticulocytes *ex vivo*. (**A**) TXB2 was fused to hFc with attenuated effector function (AEF) or fully functional wild type (WT). Mice were injected IV with 1.875 mg/kg (25 nmol/kg). Blood samples were collected after 18 hr and the percentage of reticulocytes in total number of red cells (mean ±SD, N=3) was determined by flow cytometry after staining with Thiazole Orange. (**B**) Blood samples were collected from untreated mice and the reticulocytes stained with Thiazole Orange. TXB2-hFc or 8D3-hFc antibodies were added and detected with an anti-human Fc antibody conjugated to Alexa Fluor 647. The overlaid flow cytometry histograms of reticulocytes stained with different antibodies from a single sample. (**C**) Combined data were presented as median fluorescence intensity (MFI) of antibody stained reticulocytes and was used as a measure of antibody binding (mean ±SD, N=3). Secondary antibody only staining was used as a control for background signal. * Indicates significant (p<0.05) by Student’s t-test.

## REFERENCES

1. Kawabata H. Transferrin and transferrin receptors update. Free radical biology & medicine. 2019; 133: 46–54.

2. Friden PM, Walus LR, Musso GF, Taylor MA, Malfroy B, Starzyk RM. Anti-transferrin receptor antibody and antibody-drug conjugates cross the blood-brain barrier. Proceedings of the National Academy of Sciences of the United States of America. 1991; 88: 4771–5.

3. Johnsen KB, Burkhart A, Thomsen LB, Andresen TL, Moos T. Targeting the transferrin receptor for brain drug delivery. Progress in neurobiology. 2019; 181: 101665.

4. Couch JA, Yu YJ, Zhang Y, Tarrant JM, Fuji RN, Meilandt WJ, et al. Addressing safety liabilities of TfR bispecific antibodies that cross the blood-brain barrier. Science translational medicine. 2013; 5: 183ra57, 1-12.

5. Pardridge WM, Boado RJ, Patrick DJ, Ka-Wai Hui E, Lu JZ. Blood-Brain Barrier Transport, Plasma Pharmacokinetics, and Neuropathology Following Chronic Treatment of the Rhesus Monkey with a Brain Penetrating Humanized Monoclonal Antibody Against the Human Transferrin Receptor. Molecular pharmaceutics. 2018; 15: 5207–16.

6. Lee HJ, Engelhardt B, Lesley J, Bickel U, Pardridge WM. Targeting rat anti-mouse transferrin receptor monoclonal antibodies through blood-brain barrier in mouse. The Journal of pharmacology and experimental therapeutics. 2000; 292: 1048–52.

7. Boado RJ, Zhang Y, Wang Y, Pardridge WM. Engineering and expression of a chimeric transferrin receptor monoclonal antibody for blood-brain barrier delivery in the mouse. Biotechnology and bioengineering. 2009; 102: 1251–8.

8. Yu YJ, Zhang Y, Kenrick M, Hoyte K, Luk W, Lu Y, et al. Boosting brain uptake of a therapeutic antibody by reducing its affinity for a transcytosis target. Science translational medicine. 2011; 3: 84ra44.

9. Bien-Ly N, Yu YJ, Bumbaca D, Elstrott J, Boswell CA, Zhang Y, et al. Transferrin receptor (TfR) trafficking determines brain uptake of TfR antibody affinity variants. The Journal of experimental medicine. 2014; 211: 233–44.

10. Niewoehner J, Bohrmann B, Collin L, Urich E, Sade H, Maier P, et al. Increased brain penetration and potency of a therapeutic antibody using a monovalent molecular shuttle. Neuron. 2014; 81: 49–60.

11. Karaoglu Hanzatian D, Schwartz A, Gizatullin F, Erickson J, Deng K, Villanueva R, et al. Brain uptake of multivalent and multi-specific DVD-Ig proteins after systemic administration. mAbs. 2018; 10: 765–77.

12. Webster CI, Hatcher J, Burrell M, Thom G, Thornton P, Gurrell I, et al. Enhanced delivery of IL-1 receptor antagonist to the central nervous system as a novel anti-transferrin receptor-IL-1RA fusion reverses neuropathic mechanical hypersensitivity. Pain. 2017; 158: 660–8.

13. Thom G, Burrell M, Haqqani AS, Yogi A, Lessard E, Brunette E, et al. Enhanced Delivery of Galanin Conjugates to the Brain through Bioengineering of the Anti-Transferrin Receptor Antibody OX26. Molecular pharmaceutics. 2018; 15: 1420–31.

14. Stanfield RL, Dooley H, Flajnik MF, Wilson IA. Crystal structure of a shark single-domain antibody V region in complex with lysozyme. Science (New York, NY). 2004; 305: 1770–3.

15. Griffiths K, Dolezal O, Parisi K, Angerosa J, Dogovski C, Barraclough M, et al. Shark Variable New Antigen Receptor (VNAR) Single Domain Antibody Fragments: Stability and Diagnostic Applications. Antibodies. 2013; 2: 66–81.

16. Hasler J, Rutkowski JL. Semi-synthetic nurse shark vnar libraries for making and using selective binding compounds. US20170198281A1; 2015.

17. Triguero D, Buciak J, Pardridge WM. Capillary depletion method for quantification of blood-brain barrier transport of circulating peptides and plasma proteins. Journal of neurochemistry. 1990; 54: 1882–8.

18. Lo M, Kim HS, Tong RK, Bainbridge TW, Vernes JM, Zhang Y, et al. Effector-attenuating Substitutions That Maintain Antibody Stability and Reduce Toxicity in Mice. The Journal of biological chemistry. 2017; 292: 3900–8.

19. Kissel K, Hamm S, Schulz M, Vecchi A, Garlanda C, Engelhardt B. Immunohistochemical localization of the murine transferrin receptor (TfR) on blood-tissue barriers using a novel anti-TfR monoclonal antibody. Histochemistry and cell biology. 1998; 110: 63–72.

20. Stocki P, Wicher KB, Rutkowski JL, Comper F, Demydchuk M, Szary JM. In vivo methods for selecting peptides that cross the blood brain barrier, related compositions and methods of use. 017.

21. Li L, Fang CJ, Ryan JC, Niemi EC, Lebron JA, Bjorkman PJ, et al. Binding and uptake of H-ferritin are mediated by human transferrin receptor-1. Proceedings of the National Academy of Sciences of the United States of America. 2010; 107: 3505–10.

22. Ghandour MS, Langley OK, Varga V. Immunohistological localization of gamma-glutamyltranspeptidase in cerebellum at light and electron microscope levels. Neuroscience letters. 1980; 20: 125–9.

23. Moos T. Immunohistochemical localization of intraneuronal transferrin receptor immunoreactivity in the adult mouse central nervous system. The Journal of comparative neurology. 1996; 375: 675–92.

24. Hill JM, Switzer RC, 3rd. The regional distribution and cellular localization of iron in the rat brain. Neuroscience. 1984; 11: 595–603.

25. Ghetie V, Ward ES, Vitetta ES. Pharmacokinetics of Antibodies and Immunotoxins in Mice and Humans. In: Figg WD, McLeod HL, editors. Handbook of Anticancer Pharmacokinetics and Pharmacodynamics. Totowa, NJ: Humana Press; 2004. p. 475–98.

26. Demeule M, Beaudet N, Regina A, Besserer-Offroy E, Murza A, Tetreault P, et al. Conjugation of a brain-penetrant peptide with neurotensin provides antinociceptive properties. The Journal of clinical investigation. 2014; 124: 1199–213.

27. Hermans E, Maloteaux JM. Mechanisms of regulation of neurotensin receptors. Pharmacology & therapeutics. 1998; 79: 89–104.

28. Pasqualini R, Ruoslahti E. Organ targeting in vivo using phage display peptide libraries. Nature. 1996; 380: 364–6.

29. Kolonin MG, Sun J, Do KA, Vidal CI, Ji Y, Baggerly KA, et al. Synchronous selection of homing peptides for multiple tissues by in vivo phage display. FASEB journal : official publication of the Federation of American Societies for Experimental Biology. 2006; 20: 979–81.

30. Smith MW, Al-Jayyoussi G, Gumbleton M. Peptide sequences mediating tropism to intact blood-brain barrier: an in vivo biodistribution study using phage display. Peptides. 2012; 38: 172–80.

31. Stutz CC, Georgieva JV, Shusta EV. Coupling Brain Perfusion Screens and Next Generation Sequencing to Identify Blood-Brain Barrier Binding Antibodies. AIChE journal American Institute of Chemical Engineers. 2018; 64: 4229–36.

32. Urich E, Schmucki R, Ruderisch N, Kitas E, Certa U, Jacobsen H, et al. Cargo Delivery into the Brain by in vivo identified Transport Peptides. Scientific reports. 2015; 5: 14104.

33. Li J, Feng L, Jiang X. In vivo phage display screen for peptide sequences that cross the blood-cerebrospinal-fluid barrier. Amino acids. 2015; 47: 401–5.

34. Wan XM, Chen YP, Xu WR, Yang WJ, Wen LP. Identification of nose-to-brain homing peptide through phage display. Peptides. 2009; 30: 343–50.

35. Wesolowski J, Alzogaray V, Reyelt J, Unger M, Juarez K, Urrutia M, et al. Single domain antibodies: promising experimental and therapeutic tools in infection and immunity. Medical microbiology and immunology. 2009; 198: 157–74.

36. Moos T, Morgan EH. Restricted transport of anti-transferrin receptor antibody (OX26) through the blood-brain barrier in the rat. Journal of neurochemistry. 2001; 79: 119–29.

37. Paris-Robidas S, Emond V, Tremblay C, Soulet D, Calon F. In vivo labeling of brain capillary endothelial cells after intravenous injection of monoclonal antibodies targeting the transferrin receptor. Molecular pharmacology. 2011; 80: 32–9.

38. Manich G, Cabezon I, del Valle J, Duran-Vilaregut J, Camins A, Pallas M, et al. Study of the transcytosis of an anti-transferrin receptor antibody with a Fab’ cargo across the blood-brain barrier in mice. European journal of pharmaceutical sciences : official journal of the European Federation for Pharmaceutical Sciences. 2013; 49: 556–64.

39. Sugyo A, Tsuji AB, Sudo H, Nomura F, Satoh H, Koizumi M, et al. Uptake of 111In-labeled fully human monoclonal antibody TSP-A18 reflects transferrin receptor expression in normal organs and tissues of mice. Oncology reports. 2017; 37: 1529–36.

40. Daniels-Wells TR, Penichet ML. Transferrin receptor 1: a target for antibody-mediated cancer therapy. Immunotherapy. 2016; 8: 991–4.

41. Yu YJ, Atwal JK, Zhang Y, Tong RK, Wildsmith KR, Tan C, et al. Therapeutic bispecific antibodies cross the blood-brain barrier in nonhuman primates. Science translational medicine. 2014; 6: 261ra154.

42. Sun J, Boado RJ, Pardridge WM, Sumbria RK. Plasma Pharmacokinetics of High-Affinity Transferrin Receptor Antibody-Erythropoietin Fusion Protein is a Function of Effector Attenuation in Mice. Molecular pharmaceutics. 2019; 16: 3534–43.

43. Moos T, Morgan EH. Transferrin and transferrin receptor function in brain barrier systems. Cellular and molecular neurobiology. 2000; 20: 77–95.

44. Deane R, Zheng W, Zlokovic BV. Brain capillary endothelium and choroid plexus epithelium regulate transport of transferrin-bound and free iron into the rat brain. Journal of neurochemistry. 2004; 88: 813–20.

45. Strazielle N, Ghersi-Egea JF. Potential Pathways for CNS Drug Delivery Across the Blood-Cerebrospinal Fluid Barrier. Current pharmaceutical design. 2016; 22: 5463–76.

46. Chang HY, Morrow K, Bonacquisti E, Zhang W, Shah DK. Antibody pharmacokinetics in rat brain determined using microdialysis. mAbs. 2018; 10: 843–53.

47. Mustain WC, Rychahou PG, Evers BM. The role of neurotensin in physiologic and pathologic processes. Current opinion in endocrinology, diabetes, and obesity. 2011; 18: 75–82.

48. Nemeroff CB, Bissette G, Prange AJ, Jr., Loosen PT, Barlow TS, Lipton MA. Neurotensin: central nervous system effects of a hypothalamic peptide. Brain research. 1977; 128: 485–96.

49. Popp E, Schneider A, Vogel P, Teschendorf P, Bottiger BW. Time course of the hypothermic response to continuously administered neurotensin. Neuropeptides. 2007; 41: 349–54.

50. Gu X, Wei ZZ, Espinera A, Lee JH, Ji X, Wei L, et al. Pharmacologically induced hypothermia attenuates traumatic brain injury in neonatal rats. Experimental neurology. 2015; 267: 135–42.

51. Sun YJ, Zhang ZY, Fan B, Li GY. Neuroprotection by Therapeutic Hypothermia. Frontiers in neuroscience. 2019; 13: 586.

52. Sehlin D, Pawel Stocki P, Gustavsson T, Hultqvist G, Walsh FS, Rutkowski JL, Syvänen S. Brain delivery of biologics using a cross-species reactive transferrin receptor 1 VNAR shuttle. Submitted manuscript, 2020

